# Modeling Speech-in-Noise Performance Using an Auditory-Nerve Manifold

**DOI:** 10.64898/2026.11.26.769579

**Authors:** Ahsan Jamal Cheema, Stéphane Maison, Sunil Puria

## Abstract

Speech-in-noise difficulties often exceed audiometric profile-based predictions, suggesting suprathreshold neural coding deficits limit understanding even when audibility is preserved. A major limitation of clinical speech tests is that reduced performance can reflect either hair-cell loss or cochlear neural degeneration (CND). We propose a computational framework estimating each factor’s relative contribution at a population level. We simulated auditory-nerve responses to a consonant-identification task for 94 adults (19–83 years; pure-tone average (PTA) 0–55 dB HL) under three conditions: audiometric loss alone, CND affecting low- and medium-spontaneous-rate (LS and MS) fibers, and CND affecting all fiber types. A convolutional autoencoder was used to learn low-dimensional embeddings from neurograms, and downstream classifiers were used to perform consonant identification from the embeddings. We found that audiometric loss alone, and CND of LS and MS fibers consistently produced mismatch between model and human accuracies, whereas CND affecting all fiber types predicted human accuracy closely (mean absolute error (MAE) = 0.089, *r* = 0.81, *ρ* = 0.83) on an individual level, while capturing the broad structure of human confusions.

## 1 Introduction

A central question in auditory perception research is how hearing loss and the fidelity of peripheral neural encoding shape the ability to identify speech sounds in noise. Listeners with similar hearing profiles can differ markedly in their capacity to perceive speech under adverse conditions, suggesting that standard threshold measures do not fully characterize the biological constraints on sound perception [1, 2]. Understanding which peripheral mechanisms account for this variability, and developing computational tools that can isolate their respective contributions, is essential for advancing models of auditory perception and for improving the clinical assessment of hearing function.

Clinical hearing assessment relies primarily on pure-tone thresholds to quantify audibility loss across a limited range of test frequencies [3, 4]. Threshold measures are used to derive speech intelligibility functions that generate predicted word-recognition performance in quiet [5, 6]. The magnitude of any discrepancy between predicted and measured speech recognition is clinically informative: scores that are disproportionately poor for the audiogram prompt further evaluation [7] and indicate limits on the benefit of amplification [1, 8]. At the same time, many individuals report pronounced difficulty understanding speech in noisy environments despite having normal or near-normal audiometric thresholds and word-recognition scores in quiet that align with threshold-based predictions [1, 2, 9, 10]. Large cohort studies further demonstrate considerable variability in speech-in-noise performance among listeners with matched pure-tone averages (PTAs), indicating that threshold measures do not fully capture suprathreshold distortions in temporal and spectral coding that shape real-world communication [1, 2, 5, 11].

Within the conventional framework of sensorineural hearing loss, elevated thresholds are generally interpreted as reflecting reduced cochlear amplification associated with outer hair cell (OHC) pathology [12–14], whereas speech intelligibility is thought to depend more directly on the integrity of retrocochlear speech processing [15–18]. This conceptual distinction has motivated investigation into neural mechanisms that may degrade suprathreshold performance without elevating thresholds, and cochlear neural degeneration (CND), or cochlear synaptopathy, has emerged as a candidate explanation [19]. CND reflects the loss of synapses between inner hair cells and auditory-nerve fibers (ANFs), which can degrade suprathreshold neural coding while leaving audiometric thresholds near normal [20, 21]. Animal studies show that synaptic loss can arise following noise exposure [19, 20], aging [19] or ototoxic injury [22] without measurable changes in audiometric thresholds [20, 21]. Human temporal-bone studies similarly demonstrate a significant association between age- or noise-related neural loss and poorer word-recognition performance after adjusting for hair cell loss [23–25]. Because CND reduces the number of fibers coding suprathreshold sound, particularly low-spontaneous-rate fibers whose high thresholds and wide dynamic ranges support coding at moderate-to-high levels and in noise [26, 27], it has been hypothesized to degrade speech-in-noise performance even when pure-tone thresholds remain clinically normal [20, 28]. However, translating this hypothesis into reliable clinical measures of CND has proven challenging, because CND cannot be measured directly in living listeners. Indirect noninvasive markers such as auditory brainstem responses (ABRs) [20, 29, 30], envelope-following responses [31–33] and middle-ear muscle reflexes [34–36], loudness growth [37], loss of information carrying capacity [38, 39] are linked to auditory-nerve integrity, yet findings across studies are inconsistent [40–42]. A key theme across the literature is that many physiological and behavioral measures are partly threshold-driven (via OHC amplification and thus audibility) and are also shaped by ear-canal and middle-ear acoustics, individual anatomy and central-gain factors [43, 44], which can mask or mimic synaptopathy. Interpretation is further complicated by the fact that OHC loss and CND most often co-occur and both increase with age [24, 25, 45, 46], complicating causal attribution to a specific pathology. Indeed, in a recent large-cohort study, behavioral measures designed to reveal CND, including thresholds in noise, frequency-modulation detection thresholds and speech recognition, were not predicted by otoacoustic emissions, ABRs or frequency-following responses after removing threshold-dependent variance [47]. In addition, behavioral speech-recognition measures are influenced not only by peripheral encoding fidelity but also by cognitive factors such as linguistic context, memory and attention, further complicating interpretation.

To minimize such cognitive influences, low-context speech materials are needed that remain highly sensitive to hearing impairment while reducing reliance on higher-level processing [39, 48]. The quick vowel–consonant–vowel (qVCV) test is one such speech corpus: it uses a small set of syllables embedded in speech-shaped noise at fixed signal-to-noise ratios (SNRs), and its overall accuracy and consonant-confusion patterns have been shown to separate variance related to audibility from variance that exceeds PTA-based predictions [49–51]. However, the physiological origin of this “beyond-PTA” variance remains unresolved and difficult to measure in listeners. A computational model avoids this difficulty, because audiometric loss and fiber survival can be set independently for each listener and evaluated on the same speech material [38, 52]; in such simulations, fiber loss progressively reduces the information that auditory-nerve responses carry about speech [38, 39].

Computational models have been used to study how hearing loss affects the neural coding of sound. Biophysical models of the periphery simulate hair-cell loss and the loss of low-, medium- and high-spontaneous-rate (LS, MS and HS) fibers, and relate these to evoked potentials that may serve as markers of synaptopathy [53–55]. Their consequences for speech have been examined by presenting speech processed through simulated deafferentation to normal-hearing listeners [56, 57], by predicting intelligibility from the correlation of model responses [58], by designing algorithms that restore impaired model responses [59, 60], and by decoding simulated neural responses with neural networks optimized for specific tasks [61–63]. Existing approaches, however, leave three gaps. First, the two sources of impairment have rarely been modeled together for individual listeners. Models fitted to each listener’s audiogram either omit fiber loss [62] or test only a fixed removal of LS and MS fibers [58], and where neural degeneration has been simulated for speech, it has been applied as fixed or uniform profiles, such as complete loss of LS and MS fibers or uniform loss across fiber types, that are identical for every listener [57, 59, 63]. To our knowledge, no previous model of speech perception has simulated degeneration using the age-dependent, frequency-specific fiber survival measured in human temporal bones [24], so that each listener’s neural loss follows the physiology expected for their age. Second, the decoders that link simulated neural responses to perception make strong assumptions. Networks are typically trained on large speech corpora with normal-hearing input and then tested on impaired input, so impaired responses are never seen during training [62]. The resulting errors combine the information lost in the periphery with the mismatch between impaired input and a decoder tuned to normal hearing, and the approach assumes that impaired listeners decode speech as normal-hearing listeners do; the alternative, retraining the network for each simulated periphery, assumes complete adaptation [61, 64]. Moreover, such networks are optimized to perform recognition tasks rather than to learn a compact representation of how the impaired periphery transforms the speech signal. Classifiers trained on raw neurograms depend strongly on the chosen architecture [63], and template-matching approaches rely on an ideal-observer framework with an assumed noise model, priors and similarity metric, rather than learning which features distinguish the speech sounds. In addition, to our knowledge, no previous approach has focused on (1) learning effective representations of hearing loss and neural degeneration, (2) developing models for the nonword stimuli typically used in hearing tests (such as vowel–consonant–vowel tokens, which probe phonetic cues without lexical or contextual support) and (3) learning from only the limited audio material used in a typical hearing-test session. Third, predictions are usually evaluated at the group level, so it remains unclear whether these models can track the performance of individual listeners. Autoencoders can address these gaps: rather than learning to solve a speech task, they can learn nonlinear, low-dimensional representations of high-dimensional neurograms (latent embeddings, hereafter referred to as the code, as distinct from neural coding in the auditory nerve), so that the effects of hearing loss and CND appear as changes within a latent space, and they do not require large speech corpora, making them well suited to the small stimulus sets typical of clinical speech tests.

Here, we present a stimulus-efficient framework that combines auditory-nerve modeling, autoencoder-based machine learning and human behavioral data to ask whether consonant perception in noise is better explained by hair-cell loss alone or by the addition of neural degeneration. Using the qVCV speech-in-noise test as a case study, we: (1) simulate auditory-nerve population responses to the same stimuli used behaviorally, with audiogram-matched hair-cell loss and age-scaled loss of either LS and MS fibers or all fiber types; (2) learn a low-dimensional representation of the resulting neurograms with a convolutional autoencoder shared across conditions, and identify consonants with classifiers that read this representation through listener-independent noise; and (3) compare the predicted accuracy and confusion matrices with those of 94 adults ranging from normal hearing to moderate hearing loss, each predicted by models that never saw that listener, to assess which impairment profile best reproduces individual perceptual performance. Beyond the question of CND, to our knowledge this is the first learned-representation approach developed for the nonword stimuli of a clinical hearing test. Because the autoencoder is trained without labels, the same approach could be applied to other closed-set speech tests, where it would provide a compact code of the neural response that can be interpreted directly and used to train simple downstream classifiers.

## 2 Results

### 2.1 Model Accuracy Aligns with Human Performance when Neural Degeneration is Included

We first asked how well each impairment profile predicted the overall consonant identification accuracy of individual listeners, each predicted by models that had never seen that listener’s data (Fig. 1). For each condition, the standard deviation *σ* of the decision noise was set so that the group-mean predicted accuracy matched that of the human listeners (*σ* = 0.60 for No CND and for CND of LS+MS fibers, *σ* = 0.45 for CND of all fiber types; random forest predicted accuracy 0.82, 0.80 and 0.81 vs. 0.82 for humans). CND of all fiber types gave the lowest mean absolute error (MAE) for every classifier (0.089–0.111, vs. 0.113–0.131 for No CND; Fig. 1a). Because the same 94 listeners were predicted under all three conditions, we compared conditions with paired, rank-based tests. Per-listener absolute errors are bounded at zero and strongly right-skewed (skewness 1.5–1.7; Shapiro–Wilk p < 10*^−^*^6^), so tests that assume normality are not appropriate. For each classifier, a Friedman test (a non-parametric repeated-measures analysis across the three conditions) first established an overall effect of condition. We then made the three pairwise comparisons with Wilcoxon signed-rank tests and controlled the family-wise error rate with Holm’s step-down correction, which is less conservative than Bonferroni. For the random forest, the effect of condition was significant (Friedman *χ*^2^(2) = 16.4, p < 0.001; Friedman p < 0.001 for every classifier). MAE fell from 0.113 (95% CI 0.095–0.131) without CND and 0.109 (0.094–0.125) with LS+MS CND to 0.089 (0.074–0.105) with CND of all fiber types. Prediction error was lower with CND of all fibers than with No CND in 70% of listeners (Holm-corrected p < 0.001) and lower than with LS+MS CND in 64% (p < 0.001). Both differences were significant for all four classifiers and for their average (Fig. A1). In contrast, restricting CND to LS+MS fibers did not reliably improve on No CND. The difference was not significant for the random forest (p = 0.45), XGBoost (p = 0.40), logistic regression (p = 0.68) or the average of the four classifiers (p = 0.32), and reached significance only for the SVM (p = 0.04). Predicted and actual accuracies were also most closely aligned with CND of all fiber types (Fig. 1b–d). Pearson r rose from 0.71 (No CND) to 0.75 (LS+MS) and 0.81 (all fibers), Spearman *ρ* from 0.72 to 0.79 and 0.83, and R^2^ against the identity line from 0.50 to 0.56 and 0.66. As controls, applying the identical read-out to random 16-dimensional codes yielded chance accuracy (about 10%) and no relation to listeners’ accuracy for every classifier (|r| < 0.1; Fig. A5), so the predictions arise from the learned representation of the simulated neural responses rather than from the classifiers or the decision-noise procedure. Moreover, reassigning the simulated codes at random among the held-out listeners of each fold degraded the fit in every condition and for every classifier (e.g., MAE 0.089 vs. 0.195 and r = 0.81 vs. 0.01 for CND of all fibers; p < 0.001, 10,000 permutations; Fig. A6), showing that each prediction depends on that listener’s own simulated periphery.

**Figure 1.**
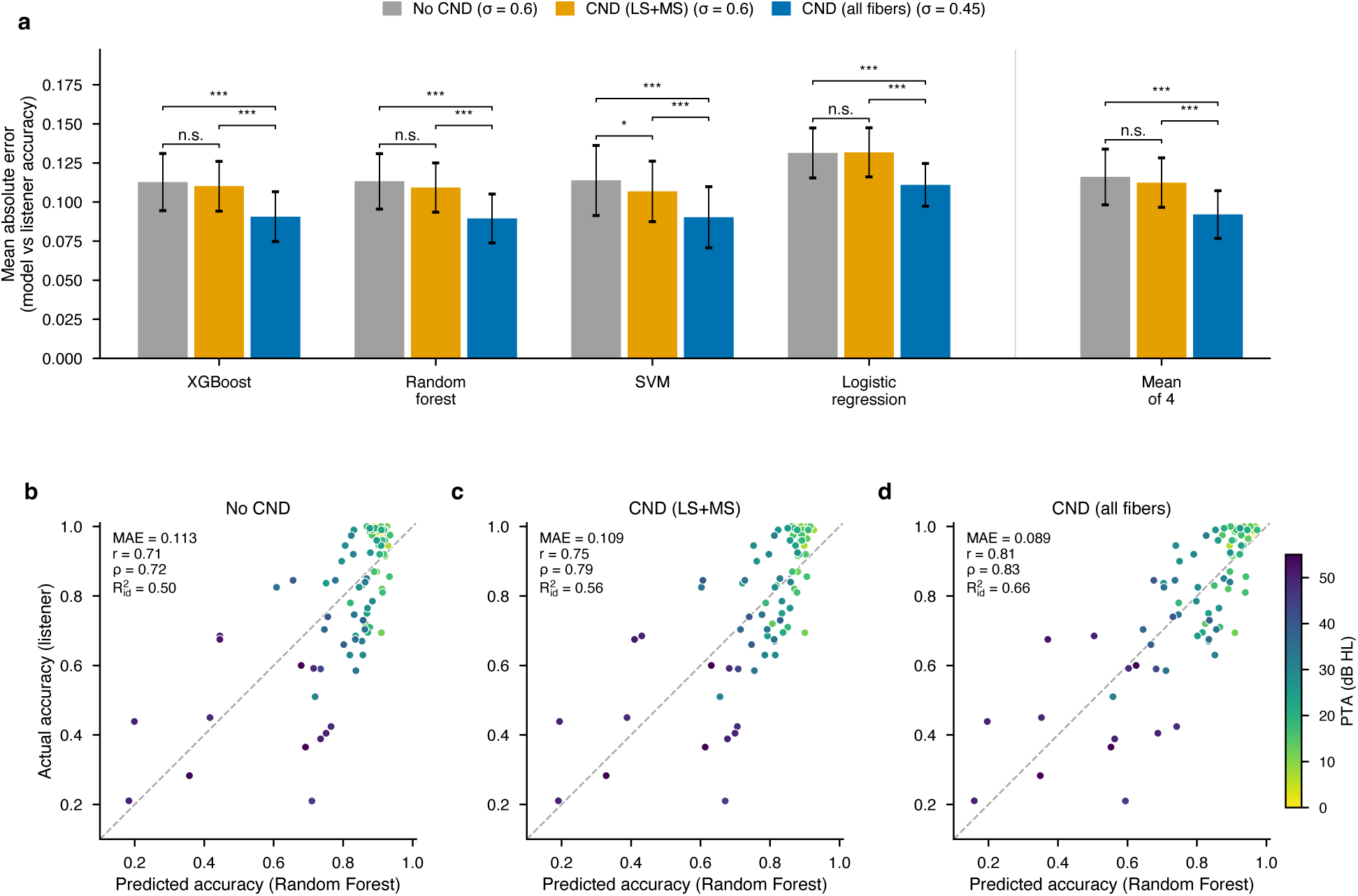
Inclusion of CND across all fiber types reduces the Mean Absolute Error between Human Subject Accuracies and Model Predicted Accuracies. Each listener’s consonant-recognition accuracy is compared with the model predicted accuracy for that listener under three simulated hearing-loss conditions. Latent embeddings from the convolutional autoencoder were used to train four downstream classifiers: XGBoost, random forest, SVM and logistic regression. Gaussian decision noise with the same standard deviation *σ* for every listener was added to the embeddings; *σ* was matched so that group-mean model accuracy equaled the human mean (*σ* = 0.60, 0.60 and 0.45). The three conditions are: audiometric loss only (No CND; outer and inner hair-cell loss), audiometric loss with age-scaled cochlear neural degeneration (CND) of low- and medium-spontaneous-rate fibers (CND LS+MS), and audiometric loss with age-scaled CND of all fiber types (CND all fibers). (a) Mean absolute error (MAE) between model and human accuracy across the 94 listeners, for each classifier and for the average of the four classifiers (“Mean of 4”). Bars show the MAE, and Error bars, 95% t-interval of the MAE across listeners. Bar color indicates the simulated condition (see legend): gray, No CND; gold, CND LS+MS; blue, CND all fibers. Brackets compare every pair of conditions using Wilcoxon signed-rank tests on the per-listener absolute errors, Holm-corrected across the three pairs (* p < 0.05; *** p < 0.001; n.s., not significant). (b–d) Predicted (random forest, the classifier with the lowest MAE) versus actual accuracy for (b) No CND, (c) CND LS+MS and (d) CND all fibers. Each point is one listener (n = 94), colored by pure-tone average (PTA, dB HL; color bar). The dashed line marks perfect prediction (predicted = actual). Each panel reports the MAE, Pearson r, Spearman *ρ* and *R*^2^_id_ (R^2^ relative to the identity line, so offsets and slope errors count against it).

The advantage of CND of all fibers does not depend on the choice of decision noise. Matching *σ* to group-mean accuracy ensures that differences between conditions reflect how well each profile orders and spaces individual listeners, not an overall offset in accuracy. Because *σ* is the read-out’s only free parameter and was fitted on the same listeners, the matched values are best-case estimates of how well each set of neurograms can predict listeners. Across the full range of *σ* (0–1; Figs. A2 and A7), no setting brought No CND or LS+MS CND to the level of CND of all fibers. Their lowest MAE (random forest: 0.111 and 0.105) and highest R^2^ against the identity line (0.51 and 0.56) remained worse than those of CND of all fibers at its matched *σ* (0.089 and 0.66), and the same held for every classifier (Figs. A2b–e and A7).

### 2.2 Model Confusions Overlap with Human Confusions and follow the Geometry of Learned Representation

Fig. 2 compares the confusions predicted by the best-performing model (random forest, CND of all fiber types) with those of the 94 listeners. Human errors were concentrated in a few confusions: place confusions between consonants sharing voicing and manner, d*→*g (15.6% of /d/ presentations), k*→*t (14.8%) and v*→*z (10.3%), and the manner confusions s*→*t (14.8%) and b*→*v (7.2%); the ten most frequent accounted for half of all errors (Fig. 2a). Model errors were concentrated instead among the fricatives and /t/, chiefly sh*→*s (11.1%), t*→*s (10.6%), s*→*t (9.3%) and s*→*z (8.8%), and were more widely spread: the ten most frequent accounted for 42% of errors, and 76 of the 90 possible confusions exceeded 0.5% of presentations, against 50 for listeners (Fig. 2b).

**Figure 2.**
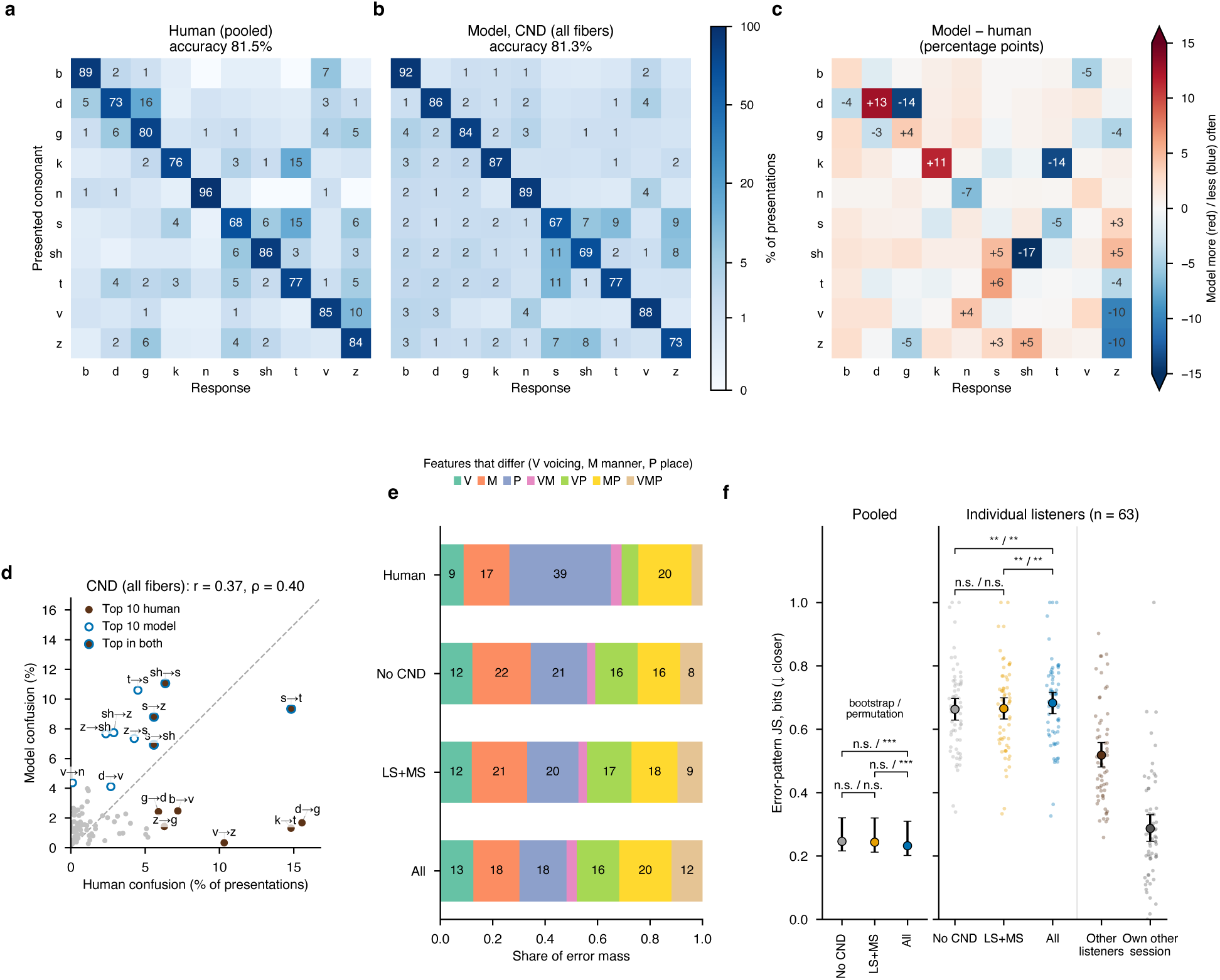
The model’s errors share the broad structure of human confusions but miss place confusions and do not capture individual listeners’ error patterns. (a) Pooled human confusion matrix: mean of the 94 row-normalized listener matrices, as % of presentations. (b) Pooled model confusion matrix for audiometric loss with age-scaled CND of all fiber types (random forest, accuracy-matched decision noise *σ* = 0.45). Both matrices are at about 81% accuracy. (c) Model minus human, in percentage points (red, the model gives this response more often than listeners; blue, less often; differences of at least 3 points are labeled). (d) Rate of each of the 90 confusions for the model versus listeners, with Pearson r and Spearman *ρ* in the title. The ten most frequent human confusions (filled brown) and model confusions (open blue) are labeled; dashed line, equal rates. (e) Error-type profile of the pooled matrices: the share of errors falling into each type, defined by which phonetic features differ between the presented consonant and the response (V, voicing; M, manner; P, place; VM, VP, MP and VMP, combinations). (f) Error-pattern Jensen–Shannon divergence (JS; off-diagonal cells renormalized to sum to 1; lower = closer) under the three model conditions. Left, pooled: JS between the pooled model matrix and the pooled human matrix of panels a–b (all 94 listeners); points, value; error bars, 95% listener-bootstrap confidence interval. Right, individual listeners: JS between each listener’s confusion matrix and the model’s prediction for that listener. Two references are shown: JS to the mean matrix of the other listeners, and JS to the listener’s own other test session (10 trials per consonant each; a conservative ceiling on how reproducible a listener’s error pattern is). Listeners with at least 5 errors and errors in both sessions (n = 63). Mean 95% listener-bootstrap confidence interval; points, individual listeners. Each bracket shows two tests between conditions, both Holm-corrected across the three pairs: a paired listener-bootstrap test (left of the slash) and a within-listener permutation test (right of the slash; condition labels swapped within each listener). Every model condition was further from the listener than the other listeners’ mean, and the other listeners’ mean was further than the listener’s own other session (paired bootstrap tests, Holm-corrected, all p < 0.001). ** p < 0.01; *** p < 0.001; n.s., not significant.

The mismatch was driven by a few consonants (Fig. 2c). The model under-predicted d*→*g (*−*13.9 percentage points), k→t (−13.5) and v→z (−10.0), and therefore identified /d/ and /k/ correctly too often, whereas it identified /sh/ and /z/ too rarely (−17.0 and −10.4 points) because it confused them with other fricatives. Across the 90 confusions, model and human rates were only moderately related (Fig. 2d; r = 0.37, *ρ* = 0.40). Four confusions were among the ten most frequent for both (s→t, sh→s, s→z, s→sh), but the other frequent human confusions were all predicted below 3%. The No CND and LS+MS models showed the same pattern (Fig. A3). Errors also differed in their phonetic features (Fig. 2e). Most human errors changed a single feature (65%), most often place alone (39%), whereas model errors changed a single feature less often (48–56%), place-only errors were half as frequent (18–21%), and voicing-plus-place or three-feature errors were more than twice as common (24–29% vs. 11%). CND of all fibers matched the human share of manner and manner-plus-place errors, but no condition reproduced the dominance of place errors.

We quantified agreement with the Jensen–Shannon divergence (JS; lower is closer) between error patterns defined as the off-diagonal cells renormalized to sum to one (Fig. 2f). At the pooled level, all models were close to the human pattern (JS = 0.246, 0.243 and 0.232 for No CND, LS+MS and all fibers). The advantage of CND of all fibers was consistent across listeners (permutation test, Holm-corrected p < 0.001) but small and not robust to resampling listeners (bootstrap, p = 0.08–0.09). For the 63 listeners with at least five errors, agreement was much weaker (JS = 0.66, 0.67 and 0.68), and CND of all fibers was slightly further from individual listeners than the other two conditions (bootstrap and permutation, p < 0.01). The other listeners’ average pattern was closer to each listener than any model (JS = 0.52), and the listener’s own other session closer still (JS = 0.29), the floor set by sampling noise. The predictions nevertheless carried some listener-specific information: the error pattern predicted for each listener was closer to that listener’s errors than the patterns predicted for the other listeners of the same fold (JS = 0.683 vs. 0.704 for CND of all fibers; p ≤ 0.002 in every condition; Fig. A6). Thus, CND of all fibers improved how many errors the model predicted for each listener (Section 2.1), but not which errors.

The source of this structure is visible in the learned representation (Fig. 3), shown as the two linear discriminants of the shared 16-dimensional code (72% of between-consonant variance). /b/, /n/, /v/ and the /k/–/g/ pair formed distinct arms, whereas /s/, /sh/, /t/ and /z/ overlapped in a central cluster with /d/ beside it (Fig. 3a–c), exactly the consonants the model confused most. Coloring by test SNR (Fig. 3d–f) shows that this cluster contains most of the tokens presented at 0 dB SNR, while consonants presented at 15–18 dB form the most distinct arms: the code is organized partly by masking, not only by phonetic features. Because the decision noise is isotropic and identical for all tokens, each token is most often misclassified as its nearest neighbors in the code (Fig. A8 shows the codes with one draw of the noise; because the noise acts in all 16 dimensions, their two-dimensional projection exaggerates the overlap between consonants). Where human confusions also fall among such neighbors, the model reproduces them: the four confusions that were frequent for both listeners and the model (s→t, sh→s, s→z and s→sh) all lie within the central cluster. The remaining human confusions (d→g, k→t, v→z, b→v) involve consonants that lie in different arms or on opposite sides of the central cluster, so the decision noise rarely carries one onto the other. Because these neighbor relations are the same in all three conditions, no impairment profile produces these confusions; the model’s errors are constrained by which clusters are adjacent in the code, and these adjacencies differ from those implied by human errors.

**Figure 3.**
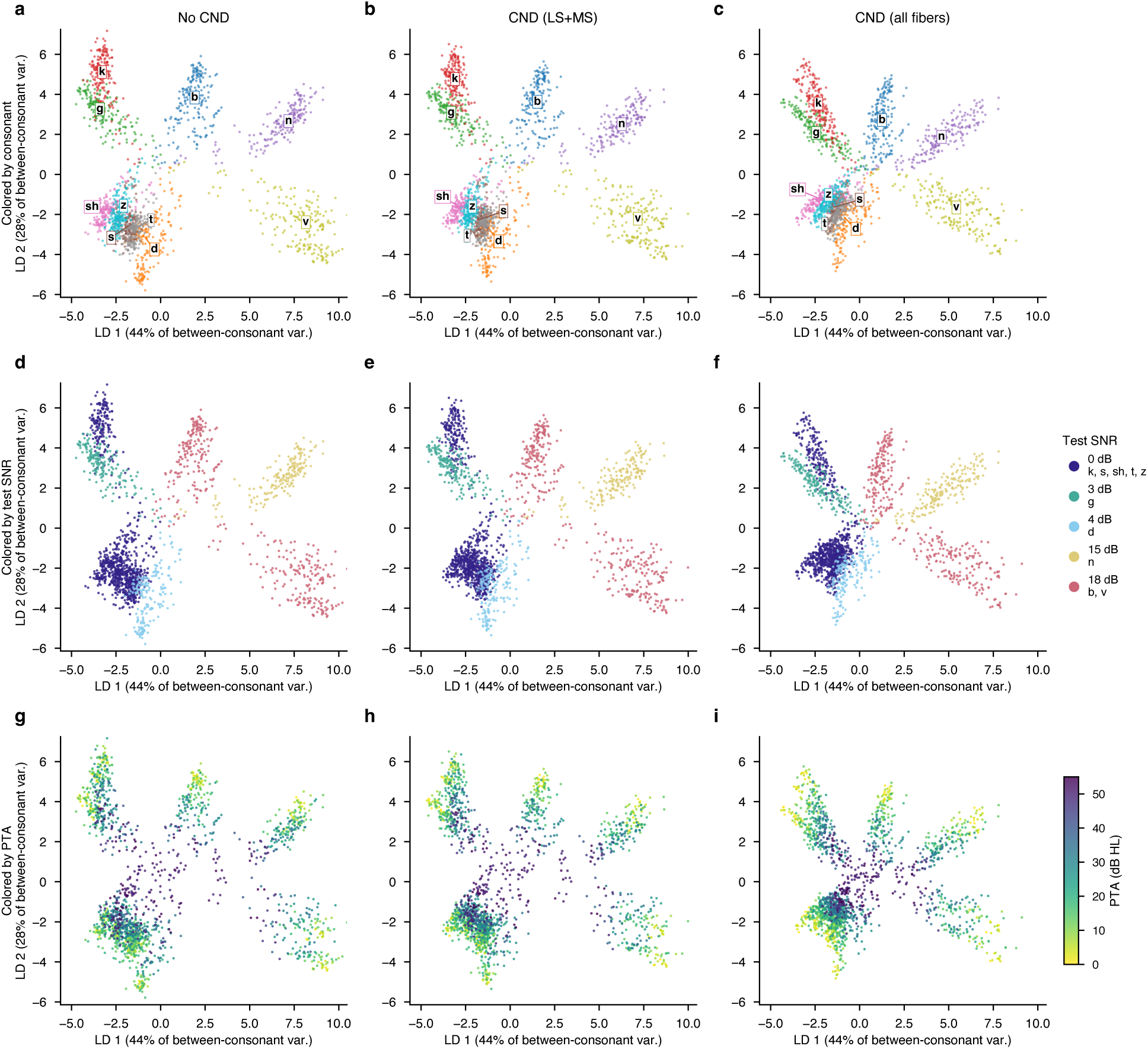
The Latent Space with hearing loss and neural degeneration. (a–c) Linear discriminant analysis (LDA) of the 16-dimensional autoencoder latent embeddings of all 94 listeners for (a) audiometric loss only (No CND), (b) audiometric loss with age-scaled CND of low- and medium-spontaneous-rate fibers (CND LS+MS) and (c) audiometric loss with age-scaled CND of all fiber types (CND all fibers). The same LDA projection was fitted on the three conditions pooled, so all panels share the same axes. Color, consonant (letters at cluster medians); axis labels give each discriminant’s share of the between-consonant variance. (d–f) The same points colored by the test signal-to-noise ratio (SNR) of each token. Each consonant was presented at a single fixed SNR (0 dB: /k/, /s/, /sh/, /t/, /z/; 3 dB: /g/; 4 dB: /d/; 15 dB: /n/; 18 dB: /b/, /v/), so SNR is confounded with consonant identity. (g–i) The same points colored by the listener’s pure-tone average (PTA, dB HL; color bar).

Coloring by PTA shows how impairment acts on this geometry (Fig. 3g–i). Codes of listeners with higher PTA lie closer to the center of each arm, where consonants are less separated, and with CND of all fibers the arms contract further. Accordingly, each listener’s total consonant separation relative to the noise tracked predicted accuracy (*ρ* = 0.91–0.95) and measured accuracy (*ρ* = 0.80–0.85; Fig. A10). Neural degeneration thus changes how far apart consonants are, and hence how many errors occur, but not which consonants are neighbors, which is set by the stimuli and their SNRs and is shared across conditions.

## 3 Discussion

The central question of this study was whether performance on a low-context consonant-in-noise test reflects audiometric loss alone or also limits in neural coding consistent with cochlear neural degeneration. Our results show that simulations incorporating age-scaled CND, particularly across all fiber types, in addition to audiogram-matched loss, more closely reproduced individual listeners’ consonant identification accuracy. The improvement held for every classifier, including a linear read-out (logistic regression) (Fig. A1), and both control analyses confirmed that it depends on each listener’s simulated periphery (Figs. A5 and A6).

This conclusion does not depend on how the decision noise was chosen. When all three conditions were read out with the same noise (*σ* = 0.45) (Figs. A4 and A9), the models without CND or with LS+MS CND overpredicted human accuracy (91% and 90% vs. 82% for listeners) and fitted individual listeners poorly (R^2^ = 0.20 and 0.29 vs. 0.66 for CND of all fibers). Matching human accuracy in the No CND and LS+MS CND conditions required adding more noise to the code (*σ* = 0.60), and even then, as at every noise level tested (Figs. A2 and A7), they fitted individual listeners worse than CND of all fibers did at lower noise (best MAE 0.111 and 0.105 vs. 0.089; best R^2^ 0.51 and 0.56 vs. 0.66). Thus, when the periphery has only audiometric loss or selective loss of LS and MS fibers, the learned representation still separates the consonants so reliably that additional variance must be added at the decision stage to bring accuracy into the human range, and even then, individual listeners are not reproduced as well. In contrast, with CND of all fibers, the noise variance required to align model and human performance is about 44% lower than in the other two conditions (*σ*^2^ = 0.20 vs. 0.36). Within this framework, degeneration across all fiber types is needed to degrade the peripheral representation enough that a modest, listener-independent decision noise yields error rates matching those of individual listeners.

The best prediction of individual listeners was obtained when CND was modeled as a distributed reduction across all spontaneous-rate (SR) fiber classes, whereas selective removal of low- and medium-SR fibers did not reliably improve on audiometric loss alone. This null result is notable given that animal models emphasize the selective vulnerability of low-SR fibers after noise exposure [27, 65]. Even though the selective vulnerability of low SR fibers in humans is difficult to measure, however human temporal-bone studies do indicate widespread age-related neural loss across the cochlea without clear evidence of SR selectivity [24, 46, 66]. Our result is also consistent with model studies in which removing low- and medium-SR fibers had little effect on predicted speech intelligibility [58] and in which speech was degraded mainly by substantial loss across fiber types [57]. This agrees with simulation studies in which speech recognition was affected only by extensive loss of auditory-nerve fibers. When speech was processed through simulated deafferentation and presented to normal-hearing listeners, recognition in noise worsened substantially only when the neural signal was very sparsely sampled [56] or when 90–95% of fibers were removed [57], and a neural-network model of speech recognition retained most of its word-recognition accuracy even after 90% loss of auditory-nerve fibers [64]. Together, these studies indicate that models typically require far more neural degeneration than selective loss of low- and medium-SR fibers to behave like listeners with impaired speech-in-noise perception. Part of the reason may be that neural-network-based decoders extract speech-relevant information from the remaining fibers very efficiently [61, 64]: because the auditory-nerve population code is highly redundant, a decoder trained on the impaired responses can recover most of the information lost with the missing fibers, whereas listeners are unlikely to decode their impaired peripheral input this efficiently. Such models therefore behave close to an ideal observer and degrade only when the loss is extensive enough to remove information that no decoder can recover. In our framework, the decision noise plays this role, limiting how precisely the code can be read out, so that a smaller, physiologically plausible loss across all fiber types is enough to bring model accuracy into the human range. In our simulations, the age-scaled loss of all fiber types removed up to 56% (apical) and 69% (basal) of fibers in the oldest listeners and, combined with audiometric loss, was sufficient to reproduce individual accuracy. We therefore interpret our results as suggesting that behaviorally relevant neural degeneration in humans may not be restricted to a single fiber class.

The low-dimensional representation also explains which confusions the model reproduces. In the shared code, most of the consonants presented at 0 dB SNR overlap in a central cluster, whereas those presented at high SNR form well-separated arms. Because the decision noise displaces each token toward its nearest neighbors, the model’s errors fall within this cluster, and the confusions that listeners also make there (s→t, sh→s, s→z and s→sh) are reproduced at the population level. The well-separated consonants remain identifiable despite the noise, and a linear read-out of the code predicted individual accuracy as well as the nonlinear classifiers (r = 0.81 for both logistic regression and random forest), indicating that the information relevant to accuracy is linearly available in the code. Hearing loss and CND act on this manifold representation by moving each listener’s codes toward the center, which reduces consonant separation and increases errors without changing which consonants are neighbors. As a result, total information coded by neurogram becomes less effective in separating the phoneme clusters and the model starts making errors that bring performance of model in human range.

Together, these findings indicate that the qVCV test captures variance related to neural coding fidelity in addition to audibility. Importantly, this conclusion arises not from correlations with physiological proxies, but from mechanistic simulations in which audiometric loss and neural survival were manipulated independently. By explicitly modeling the periphery and evaluating how different impairment profiles shape behavioral predictions, the framework provides a principled way to interpret performance in speech-in-noise tasks.

### Limitations

Several limitations should be noted. First, CND was not measured but derived from each listener’s age using published survival functions [24]. Age also covaries with central auditory processing, cognition and cumulative noise exposure, so the improvement obtained with CND of all fibers cannot be attributed specifically to peripheral neural loss alone. This shows that an age-dependent degradation of the peripheral representation, of the size expected from temporal-bone data, accounts for listeners beyond their audiograms, but other age-related factors that degrade the same information would produce a similar result. For this reason, we used a decision noise that was identical for all listeners rather than one scaled with age. Because age and hearing loss are strongly correlated in this cohort (r = 0.69 between age and PTA), age-scaled noise would have introduced a second, redundant route by which age could drive the predictions, allowing the model to track each listener’s age rather than the information in the learned low-dimensional representation. With a listener-independent noise, age influences the predictions only through the simulated neural survival. Consistent with this, predictions with CND of all fibers tracked listeners’ age slightly more strongly than their measured accuracy did (Spearman *ρ* = *−*0.82 vs. −0.74), whereas predictions without CND or with LS+MS CND tracked it less strongly (*ρ* = −0.58 and −0.68 at each condition’s own noise level, Fig. A8; −0.61 and −0.66 at a common noise level, Fig. A9); the age-scaled survival functions may therefore attribute to the periphery part of an age-related decline that has other origins. Separating peripheral from other age-related explanations will require individual estimates of neural survival that do not depend on age, for example from envelope-following or auditory brainstem responses [54, 55].

Second, the paradigm may be relatively insensitive to selective low-SR loss. Speech was presented at 65 dB SPL, where high-SR fibers are not yet saturated and still carry most of the rate information; at higher levels, where high-SR fibers saturate and the wide dynamic range of low-SR fibers becomes important, selective synaptopathy may have a larger effect.

Third, the qVCV test presents each consonant at its own fixed SNR rather than at a common or adaptive SNR, so SNR is confounded with consonant identity. The learned code is organized partly by SNR, CND has its largest effect on the low-SNR tokens, and the pattern of errors reflects the test design as well as the listener. Several frequent human confusions, d→g, k→t and v→z, were made by humans, yet the model rarely produced them under any condition, because these consonants lie in different regions of the code; they may depend on cues, such as burst spectra or formant transitions, that the code does not preserve at the resolution used here.

Relatedly, the low-dimensional manifold was learned without supervision, by reconstructing the neurograms, and is therefore not constrained to reflect the perceptual distances between phonetic categories. Its geometry is determined by the similarity of the simulated neural responses, which depends strongly on spectral shape and masking level, so consonants that listeners frequently confuse need not lie close together. This choice was deliberate, because it keeps the representation independent of the human responses it is used to predict, so that any human-like structure is a genuine prediction rather than a fitted one. Future work could learn manifolds whose distances respect phonetic structure, for example through supervised, contrastive or feature-based objectives that encode voicing, manner and place; to avoid circularity, such constraints should be derived from independent data rather than from the confusions being predicted.

Lastly, the present simulations modeled deficits at the auditory-nerve stage only. Central auditory processing was implicitly represented using downstream interpretable classifiers, however, central gain and higher-order processing may interact with peripheral degeneration, so future work extending the framework directly to brainstem and cortical stages [58] will be important for quantifying downstream consequences of CND. Crucially this extension would allow the current framework to quantify relative contribution of peripheral, neural and central stages to performance and confusions during the speech in noise test.

Despite these limitations, to our knowledge this is the first framework that uses unsupervised training to learn the hearing loss and CND embeddings rather than optimizing neural networks to solve specific tasks and it operates on the same constrained stimulus set used behaviorally, avoids reliance on large speech corpora, and remains computationally tractable and interpretable.

## 4 Conclusions

Incorporating age-scaled CND into audiogram-matched simulations improved the prediction of individual listeners’ qVCV accuracy, but only when neural loss was distributed across all fiber types; selective loss of low- and medium-SR fibers did not reliably improve on audiometric loss alone. The low-dimensional representation learned from the simulated neurograms reproduced the population-level confusions among consonants presented at low SNR, but not the place confusions that dominate the errors of listeners with hearing loss. This study introduces a data-efficient computational framework that partitions the relative contributions of audiometric loss and CND in speech-in-noise tasks, providing a mechanistic basis for interpreting suprathreshold deficits beyond the audiogram. Because the representation is learned without labels from the test’s own stimuli, the method can be applied to other closed-set hearing tests to assess efficacy of speech material, for example how well a given set of tokens and SNRs separates the predictions of different impairment profiles, and thereby to guide the design of tests that are sensitive to deficits beyond audibility loss.

## 5 Methods

### 5.1 Data Collection and Dataset Description

For this study, a total of 94 individuals ranging from normal hearing to moderately sloping hearing loss with PTA from 0 to 55 dB HL and age from 19 to 83 years old were used and their results analyzed. The clinical measurements were made at Boys Town National Research Hospital (BTNRH) and data was collected as per protocol approved by the Boys Town Institutional Review Board and data shared through a joint agreement between BTNRH and Massachusetts Eye and Ear. These individuals were presented with the VCV speech material which has been shown to be most sensitive at detecting the hearing loss [49]. This material consisted of 10 different consonants sandwiched between the same vowel /a/, each spoken by one adult female talker and recorded twice, giving 20 tokens, and sampled at 44.1 kHz. The speech level was fixed at 65 dB SPL, and stationary speech-shaped noise was added to the speech such that the SNRs were the same as those found to be most sensitive to hearing loss detection in previous studies [49–51] (Table 1). The stimuli were presented monaurally to the listeners and they were asked to identify the consonant sandwiched between the vowel /a/. The performance of the individuals was recorded in the form of confusion matrices.

**Table 1.** Phonemes Used for qVCV test and the corresponding SNRs (dB). The speech level was fixed at 65 dB SPL, and speech-shaped noise was added at the listed SNR.

| **Consonant (VCV token)** | **SNR (dB)** |
|---|---|
| aba | 18 |
| ada | 4 |
| aga | 3 |
| aka | 0 |
| ana | 15 |
| asa | 0 |
| asha | 0 |
| ata | 0 |
| ava | 18 |
| aza | 0 |

### 5.2 Auditory Nerve Model and Simulation

#### 5.2.1 Simulation of Neurograms

In this study we used an auditory nerve model (ANM) [65, 66] to simulate the responses of auditory nerve fibers (ANF) in response to the same VCV speech stimuli previously used [50, 51]. The .wav files of the speech stimuli were read using Matlab 2024 and normalized to the rms value of each wav file and converted into pressure by multiplying with the total sound pressure level (SPL) of the presentation. The wav files contained the speech signal (65 dB SPL) with additive speech-shaped noise at the SNR listed in Table 1. This pressure signal was then presented to the ANM as an input.

For this study, we used 50 cochlear frequency filters in the ANM implementation, each filter representing a unique characteristic frequency (CF) logarithmically spaced between 125 Hz – 16 kHz. Hearing loss was modeled by controlling the gains and broadness of these CF filter banks using Moore’s 2/3^rd^ outer hair cell (OHC) and 1/3^rd^ inner hair cell (IHC) loss rule [68]. For this study we used the audiometric profile of each individual listener as recorded during audiometric evaluation (Fig. 4b). These were passed to the model which then controlled the gains of OHCs and IHCs in the model. The signal from the IHCs was transduced to the ANF responses based on the model of synapse as discussed in [65, 66]. The exact number and relative proportions of different types of ANFs in humans are not known. Therefore, to model a healthy, normal-hearing cochlea, we assumed a total of about 30,000 ANFs [67] across the entire length of the cochlea. The shape of their distribution along the cochlea followed the number of ANFs per IHC in the healthy human cochlea reported by Wu et al. [24]: each of the 50 CF channels received a share of the fibers proportional to the number of ANFs per IHC at its CF multiplied by the number of IHCs it spans on the human cochlear frequency map [69], and the per-channel counts were rounded to whole fibers, so that the whole cochlea contained 29,998 fibers. Within each CF channel, 24%, 24% and 52% of the fibers were assigned to the LS, MS and HS types, respectively, as in our earlier work [52], giving 7,200 LS, 7,200 MS and 15,598 HS fibers in total. This gives the per-CF fiber distribution shown in Fig. 4d.

**Figure 4.**
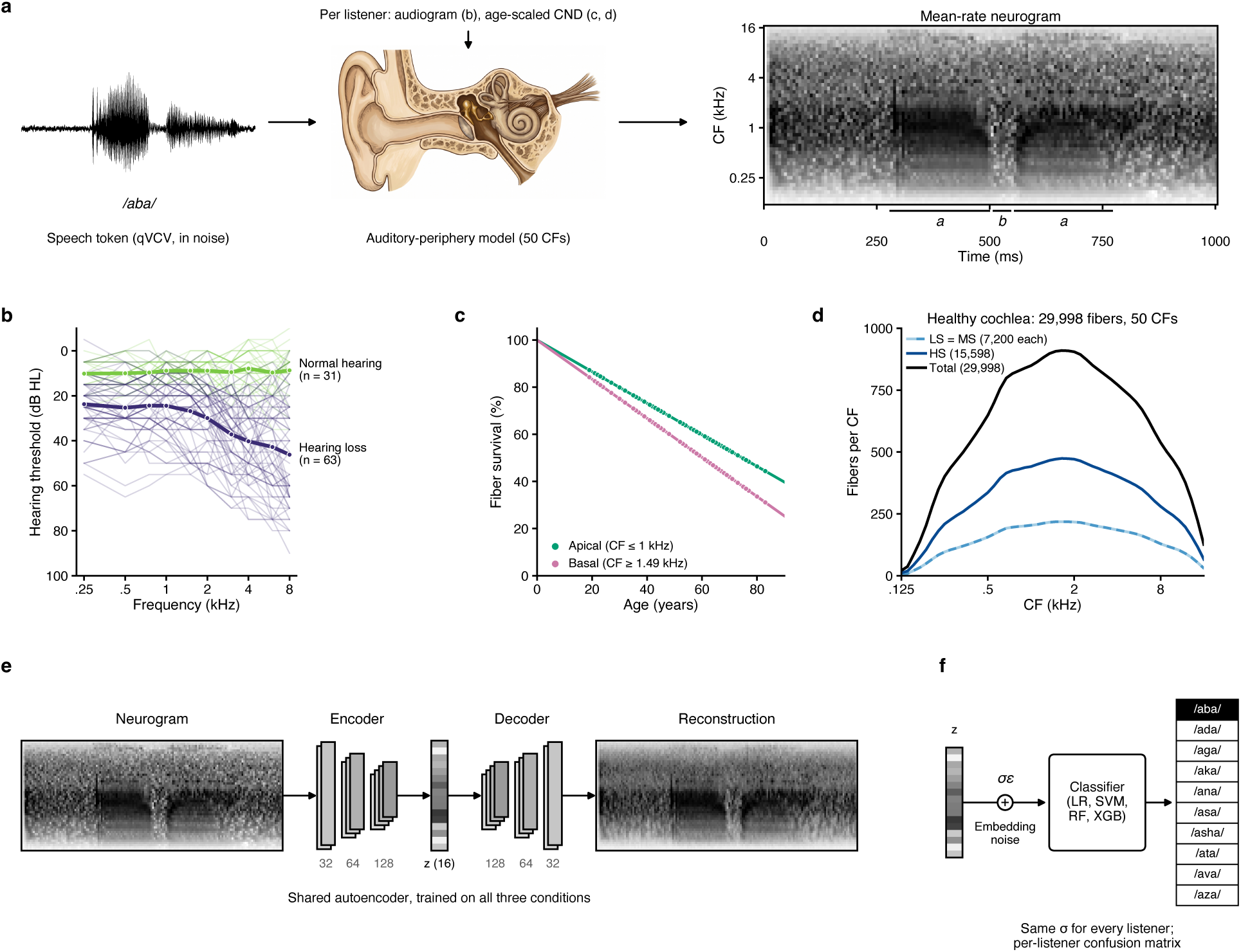
Simulation and modeling pipeline. (a) Each speech token of the quick vowel–consonant–vowel test (qVCV, presented in noise; /aba/ shown) is passed through a model of the auditory periphery with 50 characteristic frequencies (CFs), configured for each listener by their audiogram and an age-scaled loss of auditory-nerve fibers (cochlear neural degeneration, CND) to obtain a mean-rate (MR) neurogram (image of the ear and cochlea generated with Google Gemini Nano Banana Pro). (b) Audiograms of the 94 listeners (thin lines): normal hearing (n = 31) and hearing loss (slight, mild or moderate; n = 63); thick lines, group means. (c) Auditory-nerve-fiber survival as a function of age in the apical (CF 1 kHz) and basal (CF 1.49 kHz) cochlea, as used in the simulations, from Wu et al. [24]. (d) Fibers per CF in an ideal cochlea with no HL and CND, colored by spontaneous-rate type: low (LS), medium (MS) and high (HS); about 30,000 fibers in total [67], distributed along the cochlea as in Wu et al. [24]. (e) Convolutional autoencoder used for dimensionality reduction. (f) Read-out: Gaussian decision noise *σε*, with the same standard deviation *σ* for every listener, is added to z, and four classifiers identify the consonant: logistic regression (LR), support vector machine (SVM), random forest (RF) and XGBoost (XGB).

The simulations were run at a model sampling rate of 100 kHz. Synaptic outputs were represented as post-stimulus time histograms (PSTHs), which were then processed to generate neurograms: two-dimensional representations of neural activity across characteristic frequency (CF) channels and time. The mean-rate neurogram was derived by binning the PSTH into 100 *µ*s intervals and further smoothing with a 128-point Hamming window (∼12.8 ms). Responses from all fibers of a given type (LS, MS, HS) were summed across the CF axis to yield population neurograms. Together, these neurograms capture both precise temporal coding and average discharge rate across the simulated AN population, providing physiologically interpretable measures of auditory nerve activity.

For each listener, a total of 20 different Mean Rate (MR) neurograms each corresponding to the 2 recordings of the 10 /VCV/ token (Table 1) wav files presented during the experiment were simulated. In total we ended up with 1880 (2 x 10 x 94) MR neurograms.

#### 5.2.2 Simulation of Auditory Nerve Synaptopathy

To simulate CND, or auditory nerve synaptopathy, we used the ANF survival vs age data [24, 25]. For each individual’s age, the mean percentage of surviving fibers in different regions of cochlea were determined from this dataset (Fig. 4c) and incorporated into the model by reducing the proportionate number of fibers at the base of each IHC as age increased. Following the regression lines of fiber survival against age reported by Wu et al. [24], the apical (CF ≤ 1 kHz) and basal (CF ≥ 1.4 kHz) cochlea lost 0.67% and 0.83% of their fibers per year of age, respectively, with the rate interpolated on a logarithmic frequency axis in between. Since the data from Wu et al. [24, 25] only counted the number of surviving fibers and correlated them with age, and given the fact that the distribution of different fiber types in human auditory nerve has not been measured, we simulated 2 different scenarios for CND, 1) loss of fibers was assumed for all three types of fibers, and 2) loss of fibers was assumed for only LS and MS fibers, leaving the HS fibers intact. The reason for the choice of 2^nd^ scenario comes from animal studies where it has been shown that CND preferentially targets LS and MS fibers [27, 70].

### 5.3 Machine Learning Models

#### 5.3.1 Preprocessing

Since each phoneme had a different duration, the corresponding neurograms were first zero-padded so that all neurograms matched the length of the longest stimulus in the entire dataset. All neurograms spanned the same 1.19-s analysis window (187 time bins), which exceeded the duration of the longest token, so every neurogram had the same size (50 × 187). To make the input dimensions compatible with the stride-2 convolutional layers of the autoencoder, each neurogram was zero-padded to 64 CF channels (rows 51–64 set to zero) and 192 time bins, giving an input of 64 × 192; the reconstruction loss and all reported errors were computed only over the original 50 × 187 region. Neurograms were scaled by a single scalar for each cross-validation fold, equal to the maximum firing rate across the training listeners of that fold and all three conditions, so that the held-out listeners did not influence the scaling. Using one common scale for all listeners and conditions, rather than normalizing each neurogram individually, places all inputs in a range suited to the sigmoid output of the network while preserving differences in overall firing rate between listeners and conditions, including the reductions produced by hearing loss and CND, so that the learned representation reflects both the spectrotemporal pattern and the magnitude of the neural response.

#### 5.3.2 Autoencoder based model for dimensionality reduction

Convolutional autoencoders learn a compressed representation of their input by passing it through a series of convolutional layers that progressively reduce its spatial dimensions down to a low-dimensional bottleneck, and then reconstructing the input from this bottleneck with transposed convolutions. The network weights are optimized to minimize the reconstruction error between the input and its reconstruction, so that the bottleneck (the latent code) retains the features needed to reproduce the input. Because no labels are required, the representation is learned without supervision, which makes the approach suitable for small datasets such as the stimulus set of a clinical test.

The auditory-nerve simulations produced 1,880 mean-rate neurograms per condition (94 listeners × 10 consonants × 2 recordings). A single autoencoder was trained on the neurograms of all three conditions (audiometric loss only, audiometric loss with CND of LS and MS fibers, and audiometric loss with CND of all fibers), so that the three conditions share one representation and can be compared in the same coordinate system. The encoder consisted of three convolutional layers (32, 64 and 128 channels; kernel size 4, stride 2, padding 1), each followed by batch normalization and a sigmoid nonlinearity, and a linear layer that mapped the result to a 16-dimensional latent code. The latent code was batch-normalized without learnable scale or offset, so that all dimensions had zero mean and unit variance. The decoder mirrored the encoder, with a linear layer followed by transposed convolutional layers with batch normalization and sigmoid nonlinearities, and a final transposed convolution with a sigmoid output. The architecture was chosen from candidates that varied the latent dimensionality (2–16) and the number and width of the encoder layers (2–5 layers), each trained exactly as the final model, as one autoencoder for all three conditions in every cross-validation fold (Table A1). Reconstruction error fell steeply up to a 16-dimensional code and changed little beyond it (the four 16-dimensional architectures explained 92–96% of the held-out variance); among these, we used the one with the fewest parameters (1.14 million). The layers of the selected network are listed in Table A2 and the operations of every candidate in Table A3.

To avoid any leakage of listener information, all analyses used five listener-wise cross-validation folds stratified by PTA, so that each listener’s neurograms appeared in the test set of exactly one fold and were never used to train the networks or classifiers that predicted that listener. Within each fold, one autoencoder was trained on the 75–76 training listeners, of whom 15% (PTA-stratified) were held out as an inner validation set for early stopping. Networks were trained with the Adam optimizer (learning rate 10*^−^*^3^, batch size 64) to minimize the mean squared error between input and reconstruction, computed over the original 50 × 187 region of each neurogram. Training stopped when the inner-validation error had not decreased by more than 0.1% for 12 consecutive epochs (maximum 200 epochs), and the weights of the best epoch were restored. Each network was trained from three random initializations. We also trained the networks with Gaussian noise added to the latent code during training, with the same standard deviation *σ* as the decision noise used in the read-out (see below); this had little effect on the learned embeddings or on the resulting predictions, and the embeddings reported here were obtained from these networks, one set for each value of *σ*. The latent codes of the training and held-out listeners were then extracted and used to train and evaluate the consonant classifiers.

#### 5.3.3 Phoneme Recognition Classifier

Consonant identification was modeled by training classifiers to recover the presented consonant from the 16-dimensional latent code of each neurogram. To test whether the information relevant to performance is available in a simple form, we used one linear classifier (logistic regression) and three nonlinear classifiers (support vector machine with a radial basis function kernel, random forest and gradient-boosted trees, XGBoost), with standard settings that were fixed in advance and not tuned (logistic regression: L2 penalty, C = 1; support vector machine: C = 1, *γ* = “scale”; random forest: 50 trees, 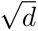 features per split; XGBoost: 150 trees, learning rate 0.05, maximum depth 6, subsampling of rows and features 0.8, L2 penalty 1), as implemented in scikit-learn and xgboost (Table A4). Classifiers were trained separately for each condition, using the same five listener-wise folds as the autoencoders, so that every listener was predicted by classifiers, and an autoencoder, that had never seen that listener’s data.

Because the latent code of a given token is deterministic, a classifier reading it directly would identify the consonants almost perfectly. To model the limited precision with which listeners make their decisions, we added Gaussian decision noise to the code before classification, in the same way during training and testing:

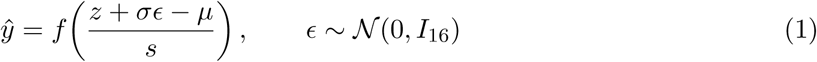

where z is the 16-dimensional latent code of a token, *ɛ* is a vector of 16 independent standard normal values (one for each latent dimension, drawn anew for every decision), *σ* is the standard deviation of the noise, *µ* and *s* are the mean and standard deviation of the noisy training codes, used to standardize the input of the classifier, and f is the classifier. Because the latent code is batch-normalized to zero mean and unit variance in each dimension, *σ* is expressed in units of the standard deviation of a code dimension: *σ* = 0.45, for example, adds noise with 45% of the spread of the codes along every dimension. The noise is isotropic and does not depend on the token, listener or condition, so it widens every consonant cluster by the same amount. The same *σ* was used for every listener, so that differences between listeners arise only from the differences in their simulated latent codes. The codes read out at a given *σ* were extracted, without noise, from the autoencoders trained with latent noise of the same *σ* (Algorithm 1).

The read-out is given step by step in Algorithm 2 and Table A4b. In each cross-validation fold, the code of every token of the training listeners was copied five times, each copy with an independent noise draw, so that each classifier learned its decision boundaries under the same noise that it would face at test. The noisy training codes were standardized with their mean and standard deviation, and the same transformation was applied to the noisy test codes. To obtain stable confusion matrices that also reflect uncertainty in the training set, each classifier was fitted 10 times, each time on a bootstrap resample of the training listeners (listeners drawn with replacement, each with all 20 of their tokens, and with new noise draws for the training copies). After each fit, every token of each held-out listener was classified twice, each time with a fresh noise draw. This yielded 400 decisions per held-out listener per classifier (10 fits *×* 2 draws × 20 tokens; 40 per consonant), which were summed into a 10 × 10 confusion matrix. Predicted accuracy was the proportion of correct decisions, and confusion matrices were row-normalized for comparison with the human data, from which “other” column responses were excluded. The procedure was repeated for three random initializations of the autoencoder, and the confusion matrices were pooled (1,200 decisions per listener and classifier).

The noise level was the only free parameter of the read-out and was set separately for each condition so that the group-mean predicted accuracy matched the group-mean human accuracy (*σ* = 0.60 for No CND and for CND of LS and MS fibers, *σ* = 0.45 for CND of all fibers), selected from a grid of 0.40–0.75 in steps of 0.05 (with 0, 0.25 and 1.0 added for the sensitivity analyses, each with one autoencoder initialization; at *σ* = 0, the autoencoder was trained without noise and the clean codes were classified directly). Because *σ* was fitted only to the group mean, the predictions for individual listeners, and their confusion patterns, were not fitted to the human data. To verify that the conclusions did not depend on this choice, all analyses were repeated across the full range of *σ* from 0 to 1 (Figs. A2 and A7).

### 5.4 Data Analysis

#### 5.4.1 Performance and comparison between model predictions and human performance

Human confusion matrices were recorded as 10 (presented) × 11 (response) counts, in which the eleventh column contained responses outside the test set (“other”). These responses were discarded, and each row was normalized to sum to one, giving a 10 × 10 matrix of response proportions for each listener. Model confusion matrices were normalized in the same way.

##### Accuracy

For each listener, predicted accuracy was the proportion of correct decisions of the model, and measured accuracy the proportion of correct responses. Agreement between predicted and measured accuracy across the 94 listeners was quantified by the mean absolute error (MAE, with a 95% t-interval across listeners), Pearson’s r and Spearman’s *ρ*, and the coefficient of determination with respect to the identity line, *R*^2^ = 1 − Σ(*y* − *ŷ*)^2^/ (*y* − *y̅*)^2^, which, unlike r, penalizes any bias or difference in scale between predicted (*ŷ*) and measured (y) accuracy. Because the same listeners were predicted under all three conditions, conditions were compared on the per-listener absolute errors with paired, rank-based tests, as these errors are bounded at zero and strongly right-skewed. For each classifier, a Friedman test assessed the overall effect of condition, and the three pairwise contrasts were tested with Wilcoxon signed-rank tests, corrected for multiple comparisons with Holm’s step-down procedure. Sign-flip permutation tests and listener-bootstrap tests of the mean paired difference gave the same pattern of results.

##### Error Pattern

To compare which confusions occur independently of overall accuracy, we defined the error pattern of a confusion matrix as its 90 off-diagonal cells, renormalized to sum to one over the whole off-diagonal. Agreement between error patterns was quantified with the Jensen–Shannon (JS) divergence,

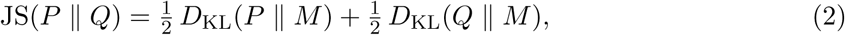

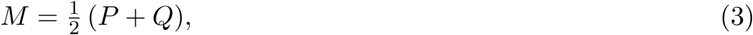

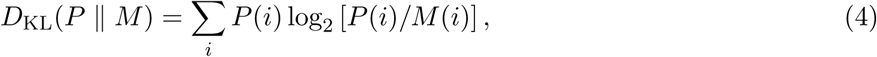

where P and Q are the human and model error patterns. JS is symmetric and bounded between 0 (identical patterns) and 1 bit (non-overlapping patterns). Because the diagonal is excluded, JS reflects the distribution of errors rather than accuracy, which was matched across conditions at the group level. Error patterns were compared at two levels. At the population level, the pooled human and model matrices were obtained by averaging the row-normalized matrices of the 94 listeners; confidence intervals were obtained by resampling listeners with replacement (5,000 resamples), and conditions were compared with a paired listener-bootstrap test and with a within-listener permutation test in which the two conditions’ matrices were swapped at random for each listener (5,000 permutations; Holm-corrected across the three pairs). At the individual level, JS was computed between each listener’s matrix and the model’s prediction for that listener, for the listeners who made at least five errors and erred in both test sessions (n = 63), because error patterns are undefined or extremely noisy for listeners with few errors. Two reference values were computed for each listener: the JS to the mean matrix of all other listeners, which indicates how well a generic population pattern predicts the listener, and the JS between the listener’s two test sessions (10 trials per consonant each), which sets a conservative ceiling on how reproducible the listener’s error pattern is. Conditions were compared with a Friedman test followed by paired listener-bootstrap and sign-flip permutation tests (Holm-corrected).

To characterize the type of errors, each confusion was classified by the phonetic features that differ between the presented and the reported consonant: voicing (voiced or voiceless), manner (stop, nasal or fricative) and place of articulation (labial, alveolar, post-alveolar or velar). The share of the pooled error mass falling into each of the seven combinations (V, M, P, VM, VP, MP, VMP) gave an error-type profile for the human and model matrices.

Consonants were classified by voicing (voiced: /b/, /d/, /g/, /n/, /v/, /z/; voiceless: /k/, /s/, /sh/, /t/), manner (stops: /b/, /d/, /g/, /k/, /t/; nasal: /n/; fricatives: /s/, /sh/, /v/, /z/) and place of articulation (labial: /b/, /v/; alveolar: /d/, /n/, /s/, /t/, /z/; post-alveolar: /sh/; velar: /g/, /k/). Each confusion was assigned to one of seven types according to which of these features differ between the presented and the reported consonant (for example, /d/→/g/ is a place error, /s/→/z/ a voicing error and /b/→/v/ a manner error).

## Statistical Testing

All statistical tests were two-sided, with *α* = 0.05. Comparisons between impairment conditions were paired, because the same listeners were predicted under all three conditions, and were performed at the level of listeners (n = 94 for accuracy; n = 63 for individual error patterns) rather than cross-validation folds. Because per-listener absolute errors and divergences are bounded and skewed, we used nonparametric tests: a Friedman test for the overall effect of condition, followed by Wilcoxon signed-rank tests for the three pairwise contrasts (accuracy), or paired listener-bootstrap tests of the mean difference (10,000 resamples) together with sign-flip permutation tests (error patterns). Population-level error patterns were compared with paired listener-bootstrap tests and within-listener permutation tests in which the conditions’ matrices were swapped at random for each listener. Within each analysis, p values were corrected across the three pairwise contrasts with Holm’s step-down procedure. Confidence intervals were 95% t-intervals across listeners (MAE) or percentile intervals from 10,000 (individual listeners) or 5,000 (pooled matrices) listener-bootstrap resamples (all other measures). No correction was applied across classifiers, because these analyses were intended to show the consistency of the effects across read-outs rather than to serve as independent tests.

## Author contributions

AJC: Conceptualization, Methodology, Software, Analysis, Original Draft, Writing and Revisions

SFM: Writing, Revisions

SP: Writing, Revisions, Data Agreements, Funding, Supervision

## Acknowledgements

This work was supported by the National Institutes of Health, National Institute on Deafness and Other Communication Disorders (NIH-NIDCD) Grant P50 DC015857 (SFM) and Massachusetts Eye and Ear, Shark Tank Research Award (SP).

The authors thank Dr. Stephen Neely for providing the dataset and arranging the data-sharing agreement.

## Competing interests

The authors declare no competing interests.

## Appendix

## A Appendix figures

**Figure A1.**
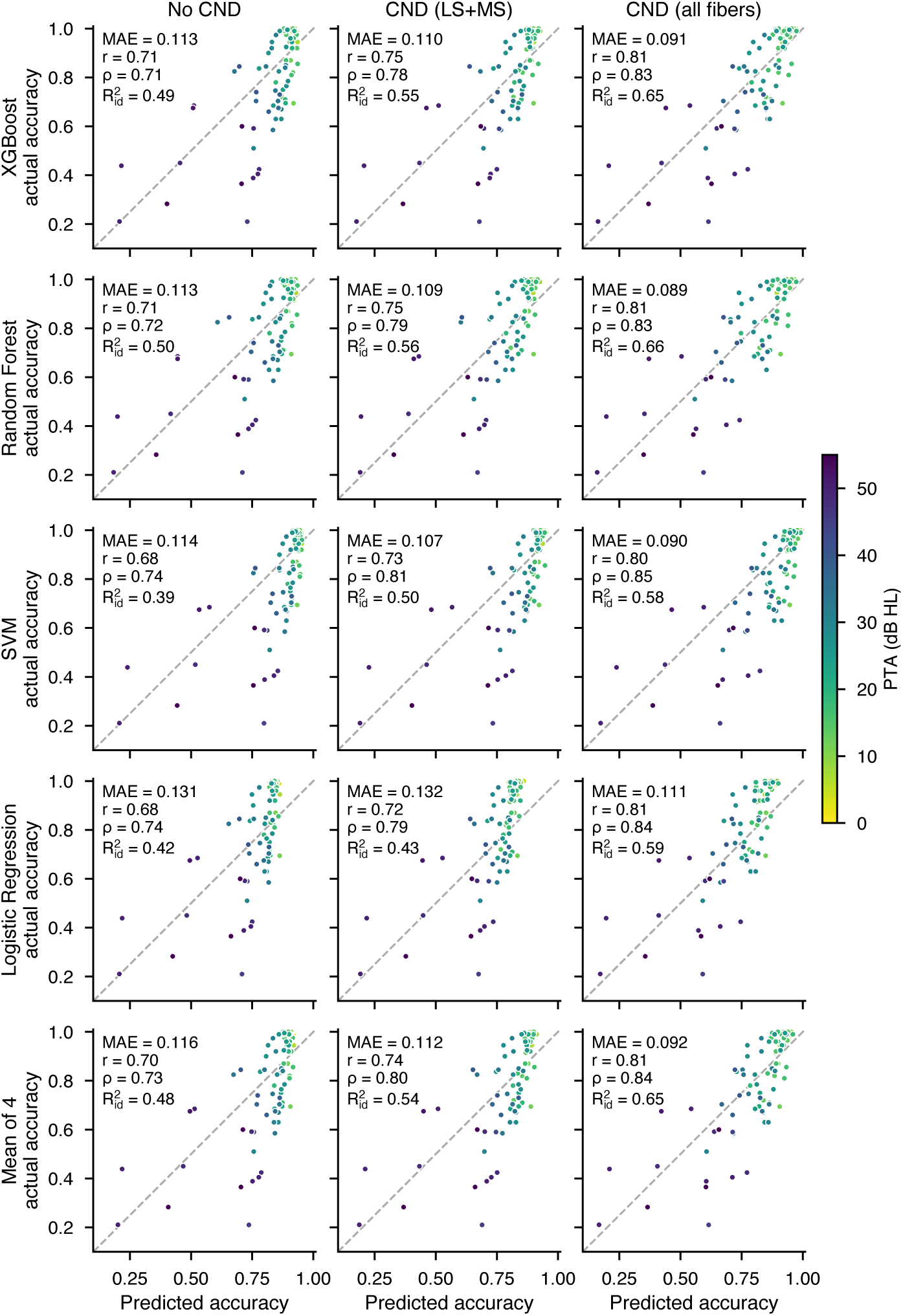
CND of all fiber types gives the closest prediction of individual accuracy for every classifier. Predicted versus actual accuracy for every classifier. Each panel plots the measured consonant identification accuracy of each of the 94 listeners against the accuracy predicted by models that never saw that listener (five listener-wise cross-validation folds), for the three impairment conditions (columns: audiometric loss only, No CND; with CND of low- and medium-spontaneous-rate fibers, CND (LS+MS); and with CND of all fiber types, CND (all fibers)) and each read-out (rows: XGBoost, random forest, support vector machine, logistic regression, and the mean of the four classifiers, obtained by pooling their predicted confusion matrices). Decision noise was set separately for each condition to match the group-mean human accuracy (*σ* = 0.60, 0.60 and 0.45), with the same value for all classifiers. Points are listeners, colored by PTA; the dashed line marks equal predicted and actual accuracy. Each panel reports the mean absolute error (MAE), Pearson r, Spearman *ρ* and the coefficient of determination with respect to the identity line (R^2^id). For every classifier, CND of all fiber types gave the lowest MAE and the highest r and R^2^id, including for the linear read-out (logistic regression).

**Figure A2a.**
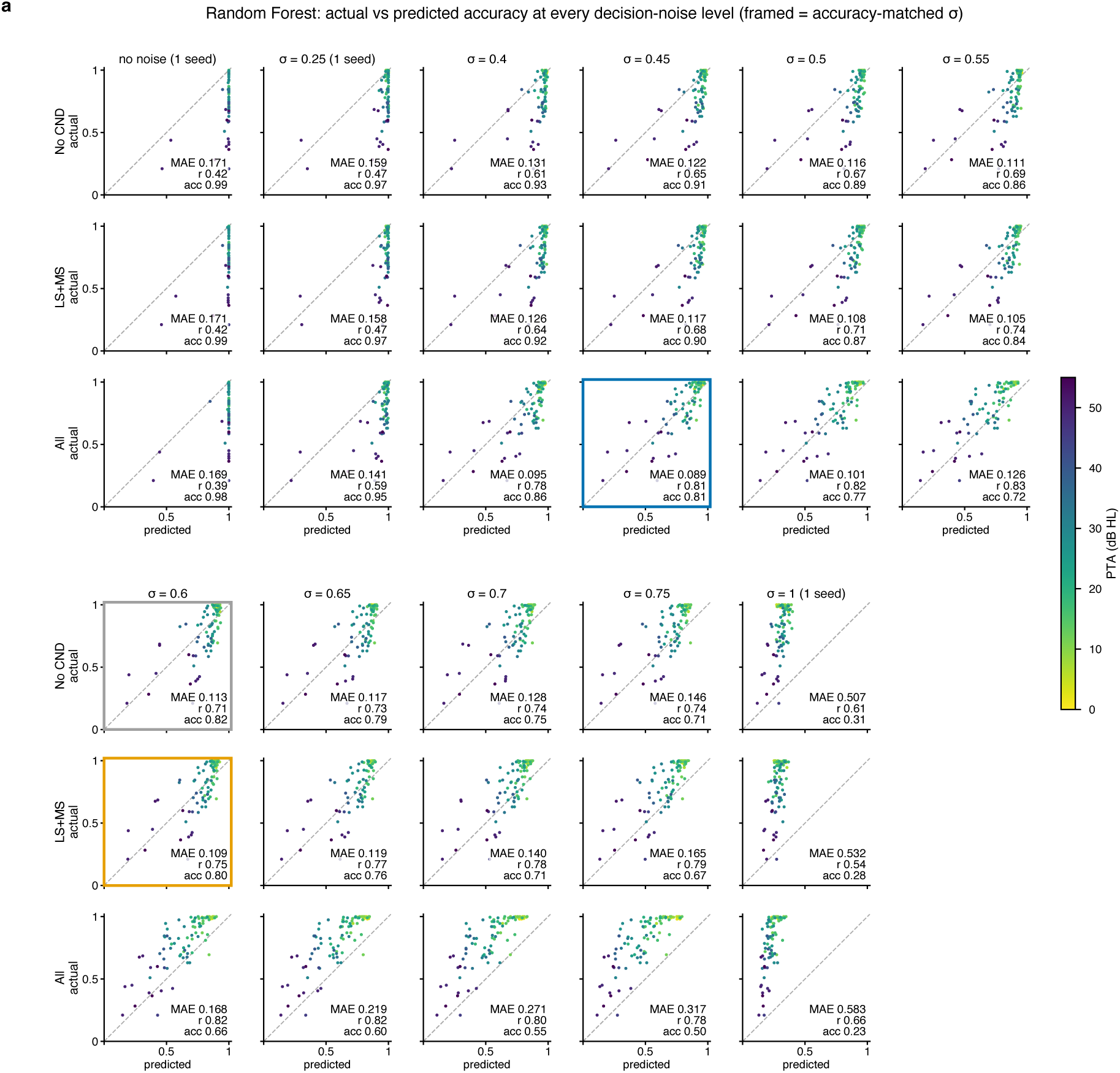
No level of decision noise brings audiometric loss alone or CND of LS and MS fibers to the agreement obtained with CND of all fibers. Predicted versus actual accuracy across decision-noise levels (random forest). Each panel plots the measured accuracy of each of the 94 listeners against the accuracy predicted by the random forest read-out, for one impairment condition (rows: audiometric loss only, No CND; with CND of low- and medium-spontaneous-rate fibers, LS+MS; with CND of all fiber types, All) and one level of decision noise (columns: *σ* = 0, i.e. a noise-free read-out, 0.25, 0.40–0.75 in steps of 0.05, and 1.0; the grid continues in the lower block). Points are listeners, colored by PTA; the dashed line marks equal predicted and actual accuracy. Each panel reports the mean absolute error (MAE), Pearson r and the group-mean predicted accuracy (acc; human group mean = 0.82). Framed panels mark the accuracy-matched noise level of each condition used in the main analyses (gray, No CND, *σ* = 0.60; orange, LS+MS, *σ* = 0.60; blue, All, *σ* = 0.45). Results for *σ* = 0, 0.25 and 1.0 are from one autoencoder initialization; all other levels pool three initializations. Without decision noise, every condition predicted near-perfect accuracy for almost all listeners; increasing *σ* lowered predicted accuracy and spread the predictions. The lowest MAE was reached at *σ* = 0.55 for No CND (0.111) and LS+MS (0.105), and at *σ* = 0.45 for CND of all fibers (0.089); no noise level brought No CND or LS+MS to the agreement obtained with CND of all fibers.

**Figure A2b.**
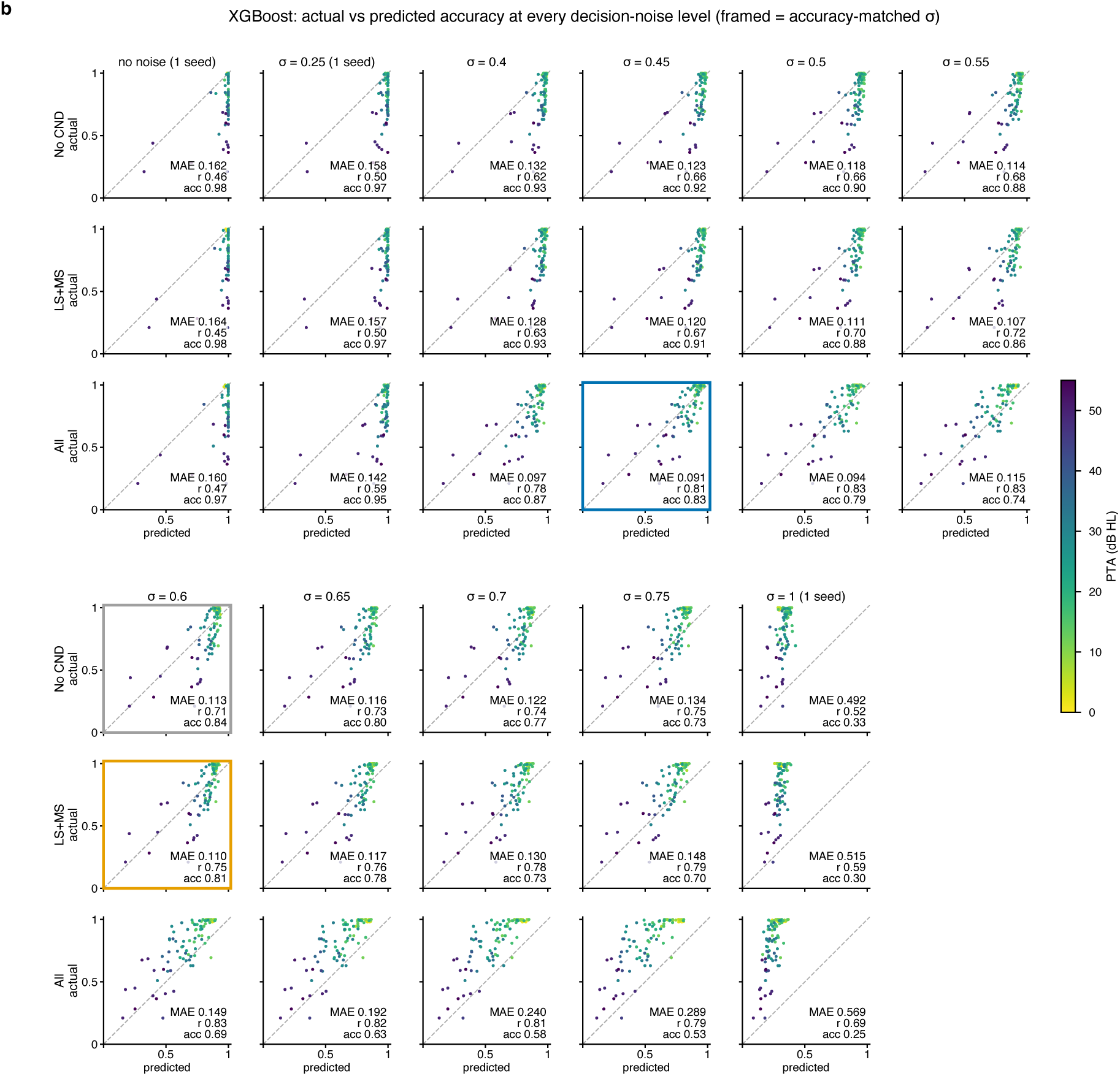
As Fig. A2a, for XGBoost.

**Figure A2c.**
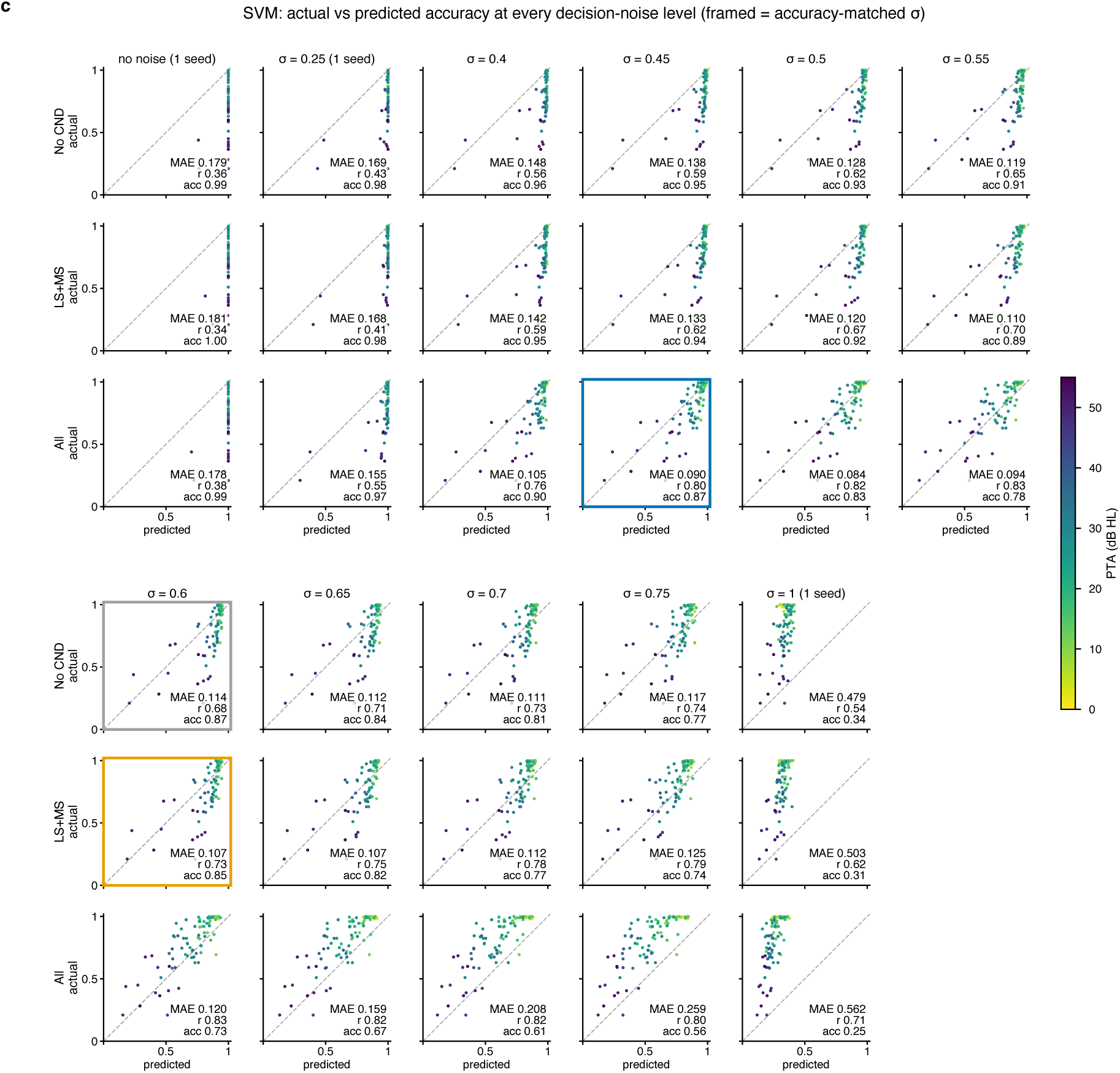
As Fig. A2a, for the support vector machine (SVM).

**Figure A2d.**
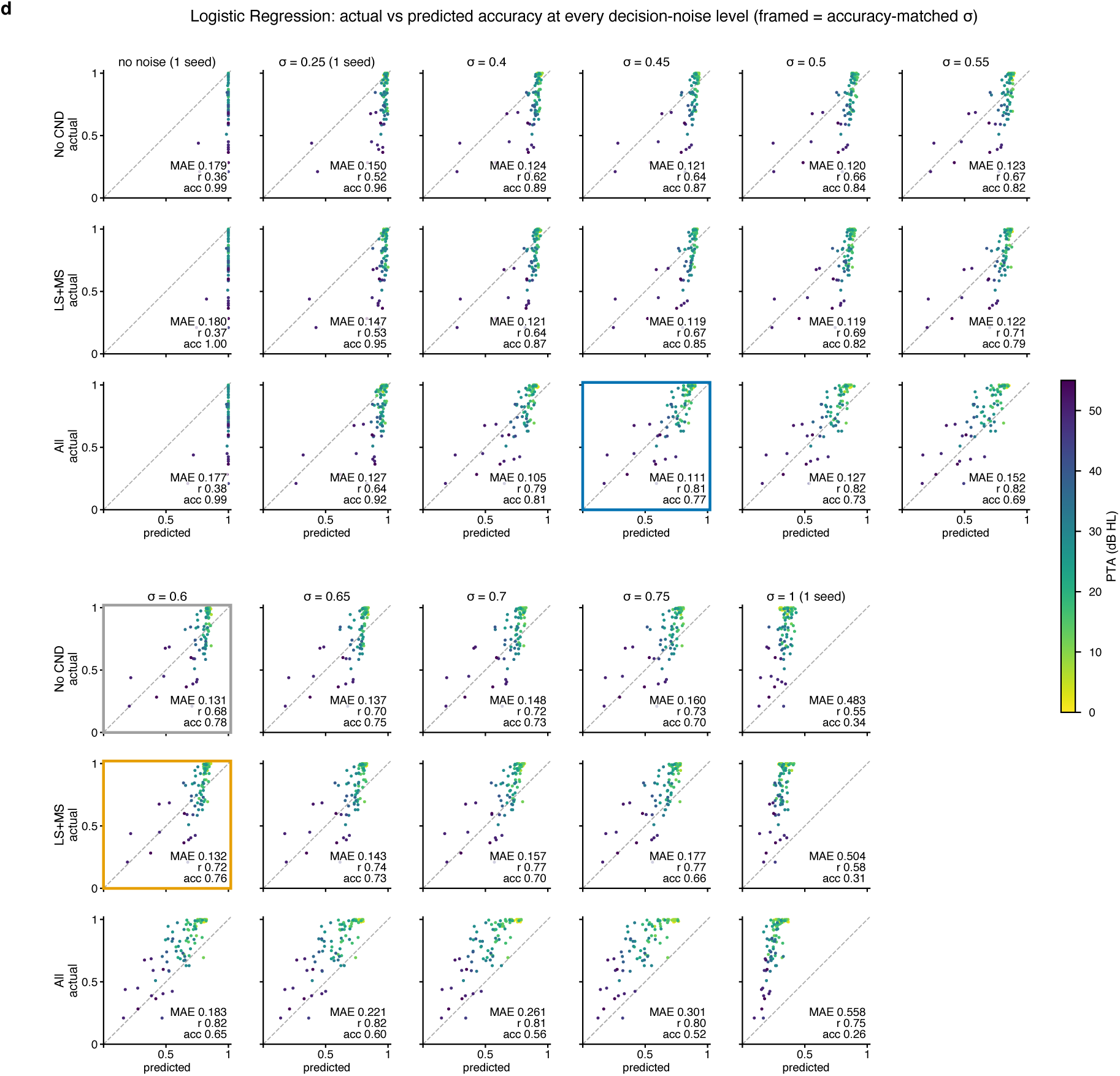
As Fig. A2a, for logistic regression.

**Figure A2e.**
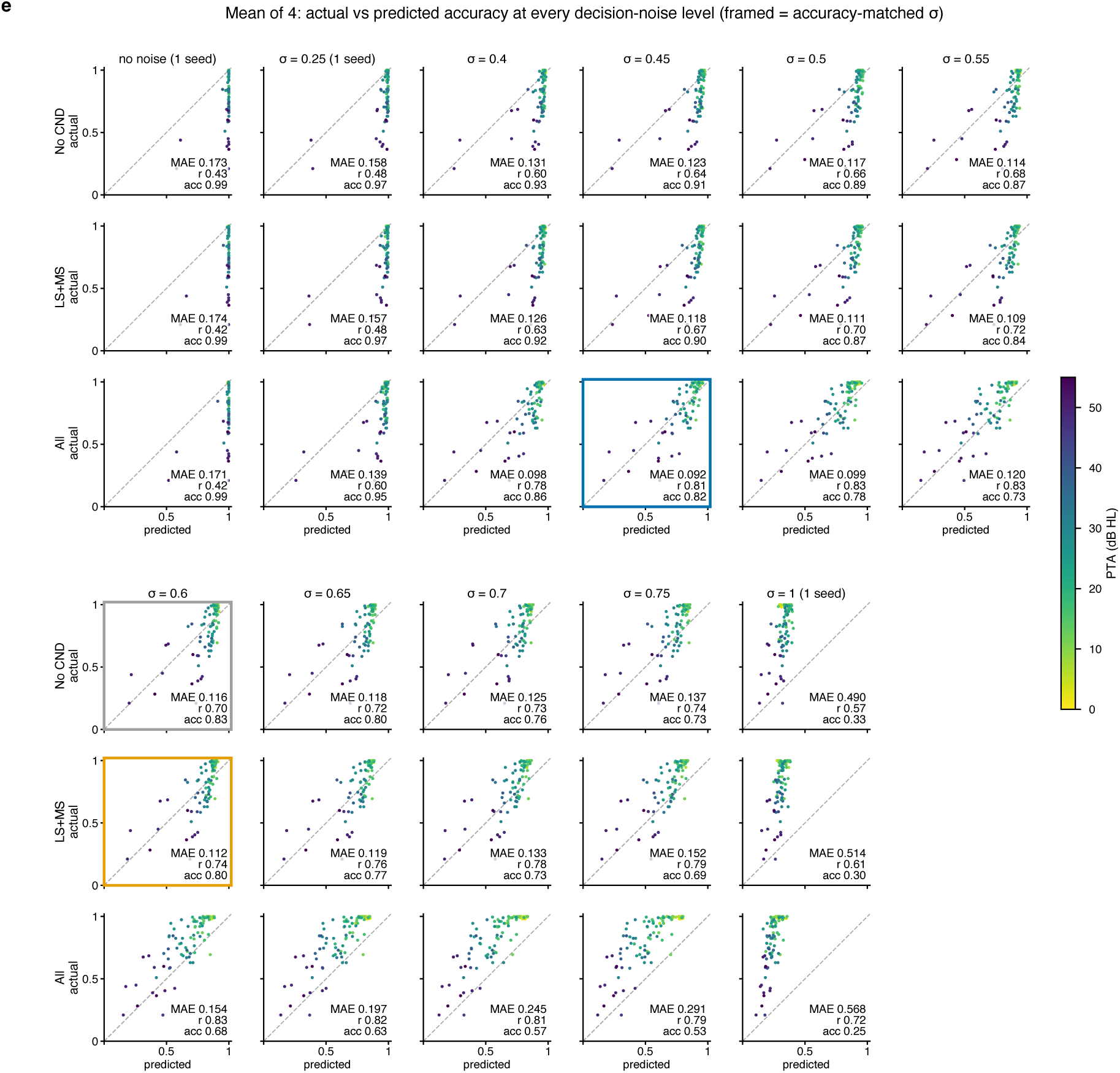
As Fig. A2a, for the mean of the four classifiers.

**Figure A3.**
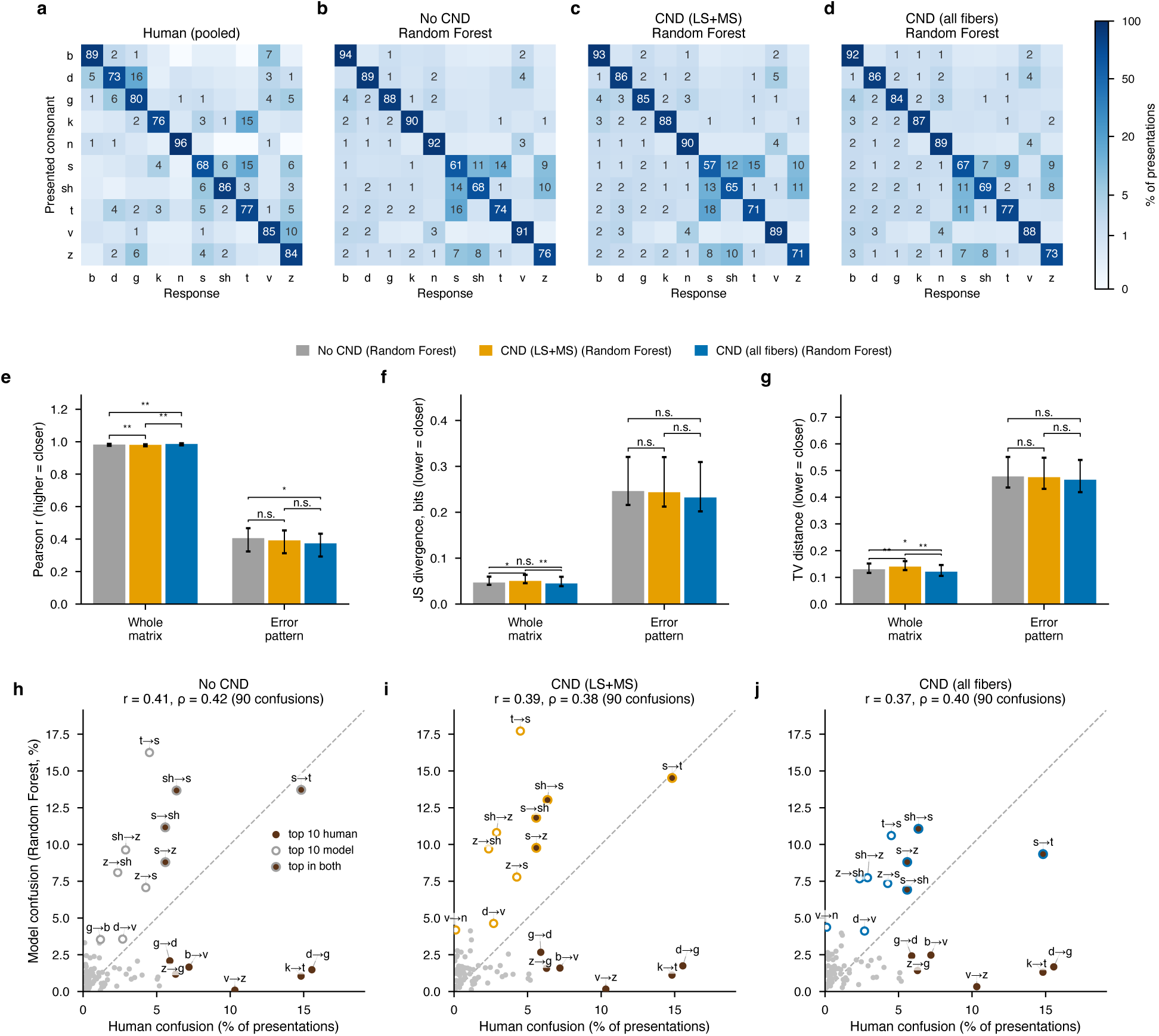
All three conditions produce the same pattern of confusions, concentrated among /s/, /sh/, /t/ and /z/, and miss listeners’ place confusions. Pooled confusion matrices of all three impairment conditions. (a) Pooled human confusion matrix: mean of the 94 row-normalized listener matrices, as % of presentations (“other” responses excluded). (b–d) Pooled confusion matrices predicted for the same listeners by the random forest read-out with accuracy-matched decision noise, for audiometric loss only (b, No CND, *σ* = 0.60), with CND of low- and medium-spontaneous-rate fibers (c, CND (LS+MS), *σ* = 0.60) and with CND of all fiber types (d, CND (all fibers), *σ* = 0.45). (e–g) Agreement between each pooled model matrix and the pooled human matrix, for the whole matrix (left) and for the error pattern, i.e. the 90 off-diagonal cells renormalized to sum to one (right), measured by Pearson r (e; higher = closer), Jensen–Shannon divergence (f; bits, lower = closer) and total-variation distance (g; lower = closer). Bars, values for the pooled matrices; error bars, 95% percentile intervals from resampling listeners with replacement (5,000 resamples). Brackets, paired listener-bootstrap tests between conditions, Holm-corrected across the three pairs (* p < 0.05; ** p < 0.01; n.s., not significant). (h–j) Rate of each of the 90 confusions for the model versus listeners in each condition, with Pearson r and Spearman *ρ*; the ten most frequent human confusions (filled brown) and model confusions (open, condition color) are labeled; dashed line, equal rates. In all three conditions the model’s most frequent confusions are among /s/, /sh/, /t/ and /z/, and the frequent human place confusions (d→g, k→t, v→z) are rarely produced.

**Figure A4.**
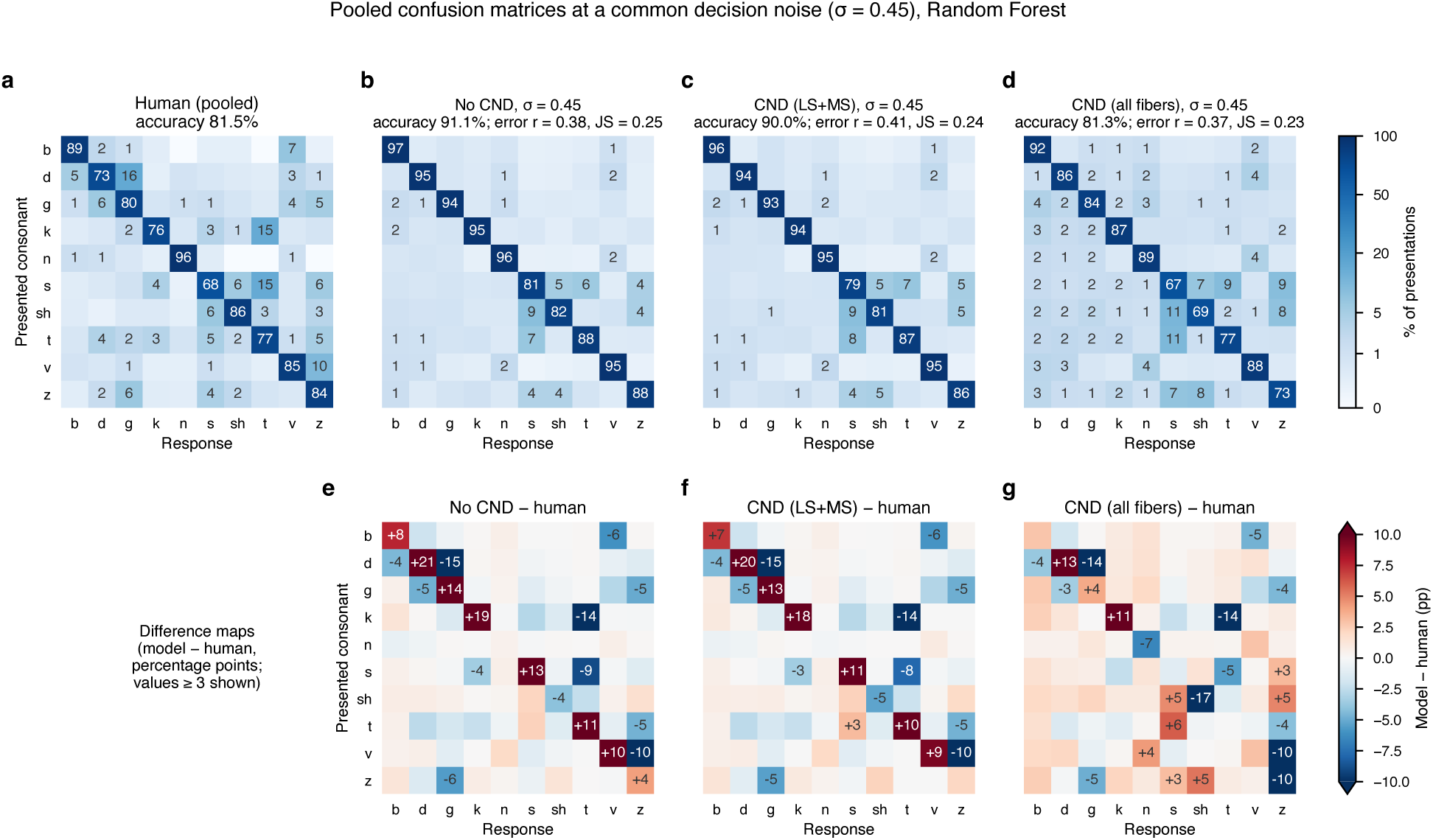
At a common decision noise, only CND of all fibers lowers accuracy to the human level, while the pattern of errors stays the same. Pooled confusion matrices of all three conditions at a common decision noise. (a) Pooled human confusion matrix (mean of the 94 row-normalized listener matrices, % of presentations; accuracy 81.5%). (b–d) Pooled confusion matrices predicted by the random forest read-out when all three conditions are read out with the same decision noise (*σ* = 0.45, the accuracy-matched value for CND of all fiber types), for audiometric loss only (b, No CND), with CND of low- and medium-spontaneous-rate fibers (c, CND (LS+MS)) and with CND of all fiber types (d, CND (all fibers)). Titles give the overall accuracy and the agreement of the error pattern (the 90 off-diagonal cells renormalized to sum to one) with the human pattern, as Pearson r and Jensen–Shannon divergence (JS, bits). (e–g) Model minus human, in percentage points (red, the model gives this response more often than listeners; blue, less often; differences of at least 3 points are labeled). At the same decision noise, the models without CND or with LS+MS CND identify almost every consonant too often (diagonal entries 79–97%; overall accuracy 91.1% and 90.0% vs. 81.5% for listeners), whereas CND of all fibers matches human accuracy (81.3%). The pattern of errors is similar in all three conditions and differs from the human pattern in the same way, with too few place confusions (d→g, k→t, v→z) and more confusions among /s/, /sh/, /t/ and /z/.

**Figure A5.**
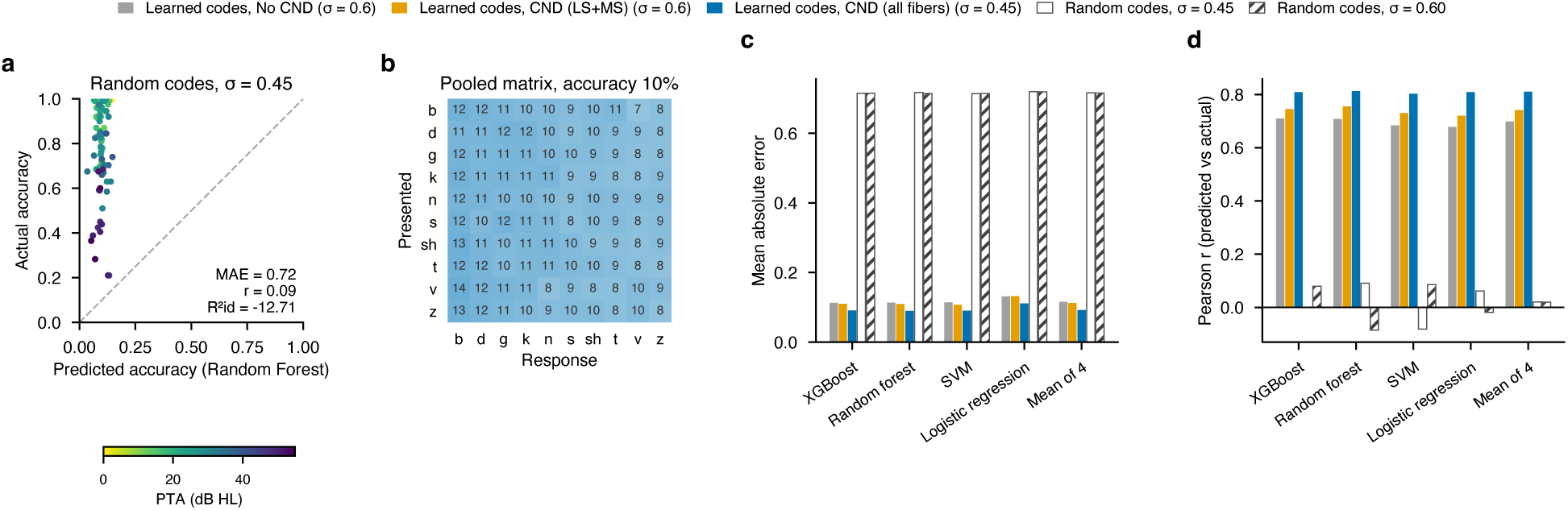
Random codes give chance accuracy and no relation to listeners, so the predictions come from the learned representation. Control: the read-out applied to random codes. Each of the 1,880 tokens (94 listeners 10 consonants 2 recordings) was assigned an independent random 16-dimensional code drawn from N(0, I), matching the scale of the batch-normalized learned codes, and the complete read-out of the main analysis was repeated unchanged: the same five cross-validation folds, the same four classifiers and settings, five noisy copies per training token, and 10 listener-bootstrap fits with two noisy draws per held-out token. Random codes contain no information about the stimulus, hearing loss or fiber type, so a single control applies to all three conditions. (a) Measured versus predicted accuracy of each listener for the random forest with random codes at *σ* = 0.45; points colored by PTA; dashed line, equal predicted and actual accuracy. (b) Pooled predicted confusion matrix for the same read-out (% of presentations). (c, d) Mean absolute error (c) and Pearson correlation between predicted and measured accuracy (d) for each classifier, for the learned codes of each condition at its accuracy-matched decision noise (gray, No CND, *σ* = 0.60; orange, CND (LS+MS), *σ* = 0.60; blue, CND (all fibers), *σ* = 0.45) and for the random codes at *σ* = 0.45 (open bars) and *σ* = 0.60 (hatched bars); results without decision noise were the same. With random codes, every classifier performed at chance (accuracy 10%) with uniform confusions, and predicted accuracy was unrelated to listeners’ accuracy (r between 0.09 and 0.09), whereas the learned codes of all three conditions predicted individual accuracy, most closely with CND of all fiber types. The predictions therefore arise from the learned representation of the simulated neural responses, not from the classifiers or the decision-noise procedure. The decision-noise procedure is required to calibrate the separability of the consonant codes.

**Figure A6.**
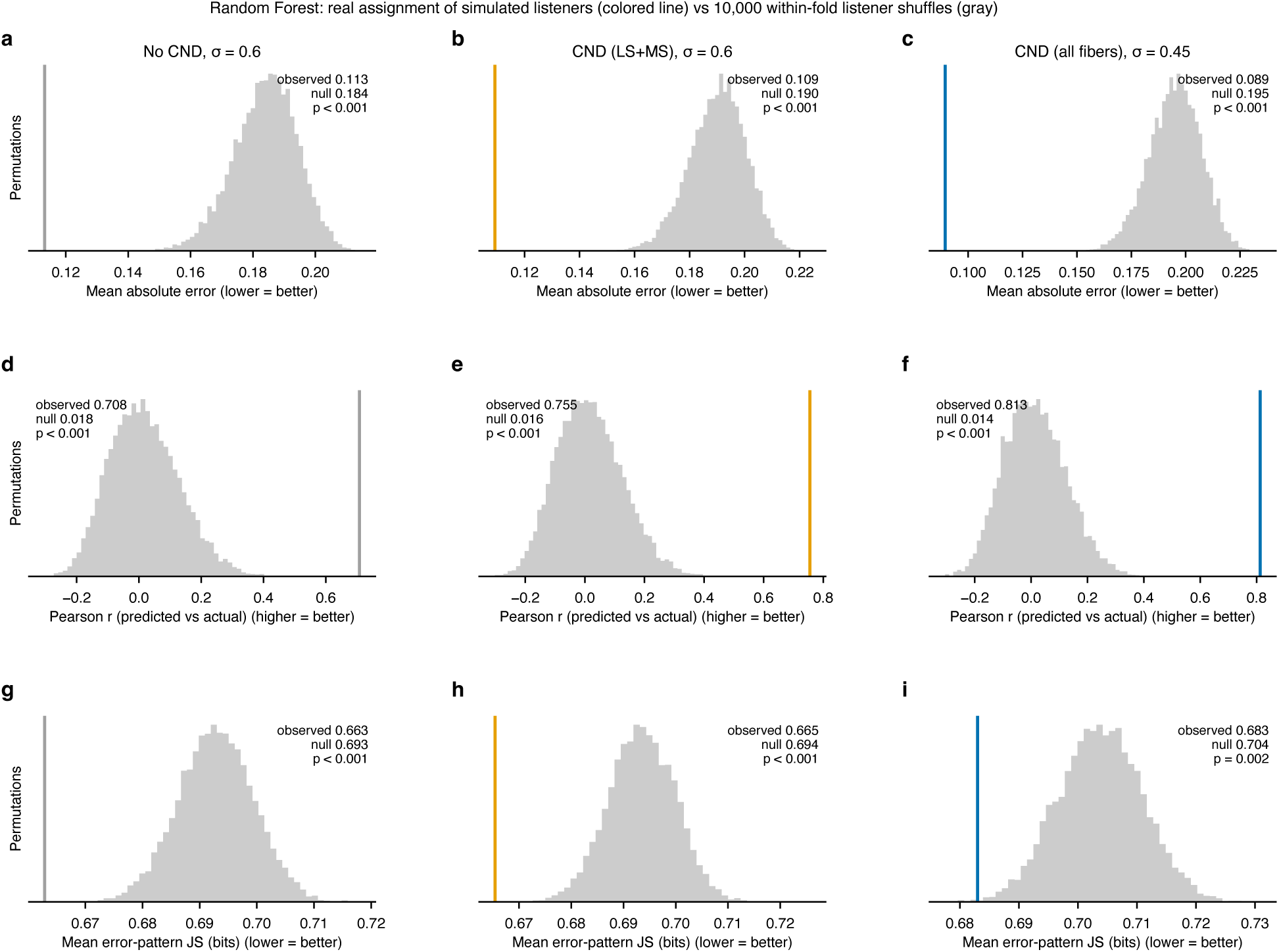
Predictions depend on each listener’s own simulated periphery: reassigning codes among listeners degrades accuracy and error-pattern agreement. Listener-shuffle permutation control. Within each cross-validation fold, the simulated codes of the held-out listeners were reassigned at random to the other held-out listeners of the same fold (10,000 permutations; folds are PTA-stratified, so every fold spans the full range of hearing). Because the classifiers of a fold are fixed, this is equivalent to repeating the read-out with each listener’s simulated periphery replaced by that of another listener. Histograms show the null distributions for the random forest read-out at the accuracy-matched decision noise of each condition (columns), and colored lines the values for the real assignment: mean absolute error between predicted and measured accuracy (a–c; all 94 listeners), Pearson correlation between predicted and measured accuracy (d–f) and mean error-pattern Jensen–Shannon divergence between each listener and the prediction made for that listener (g–i; listeners with at least five errors and errors in both sessions, n = 63). The key comparison is between each colored line and the gray distribution. If the predictions did not depend on each listener’s own simulated codes, any listener’s codes would predict a given listener equally well, and the colored line would fall within the gray distribution. A colored line lying outside the distribution, to the left for MAE and JS (lower is better) and to the right for r (higher is better), shows that the correct assignment predicts better than random reassignments. For accuracy (a–f), the lines lie far outside the distributions in every condition, whereas for error patterns (g–i) they lie just beyond the lower edge, so the listener-specific information about which errors are made is real but small. One-sided p values are the proportion of permutations at least as good as the real assignment. For every condition and classifier, the real assignment outperformed all permutations for accuracy (p < 0.001), and the predicted error patterns were closer to the corresponding listener than to other listeners (p 0.002), showing that the predictions depend on each listener’s own simulated periphery.

**Figure A7.**
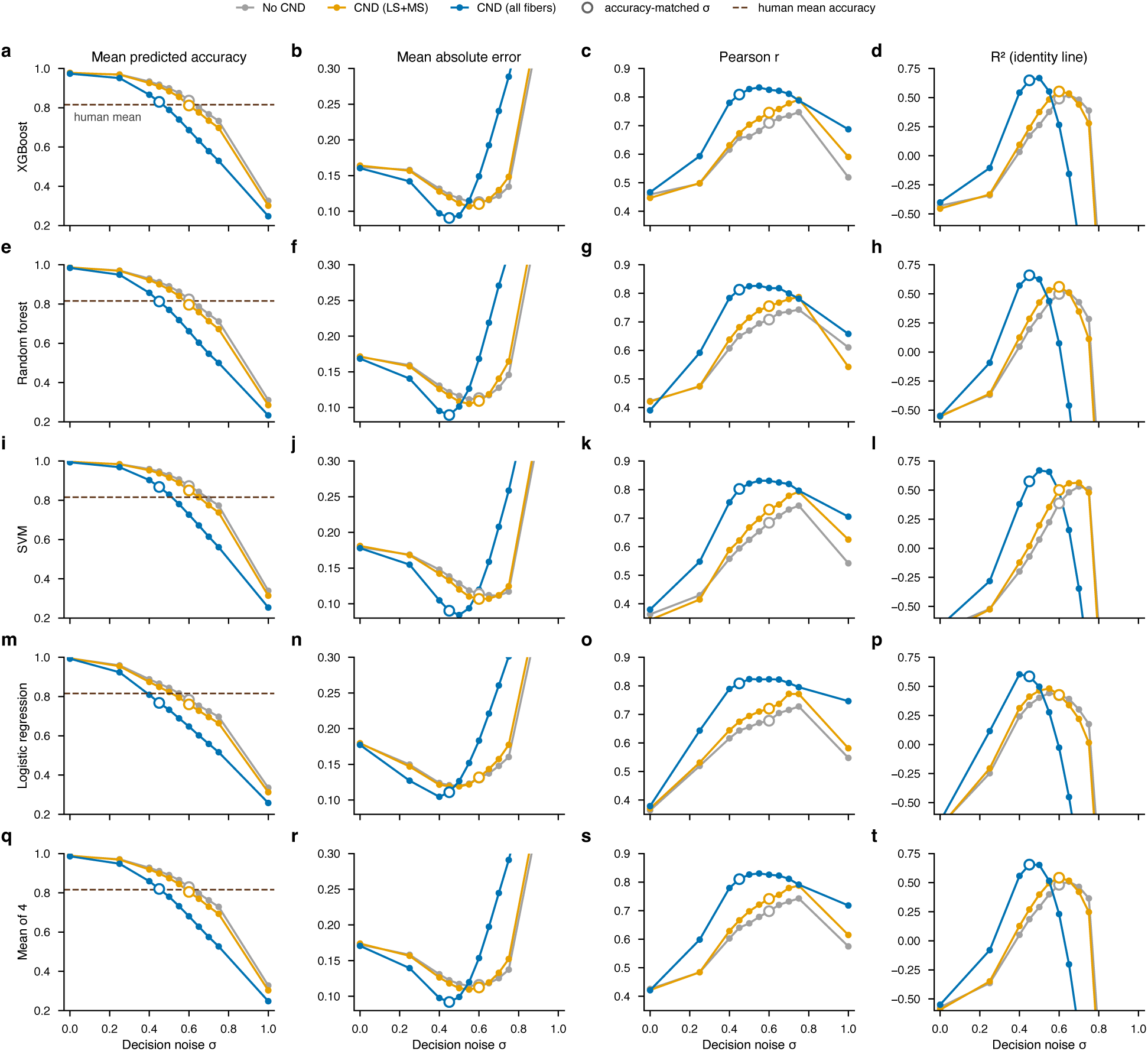
For every read-out, no decision-noise level lets audiometric loss alone or CND of LS and MS fibers match the agreement of CND of all fibers. Agreement with listeners across decision-noise levels. For each read-out (rows) and impairment condition (colors), the group-mean predicted accuracy (first column; dashed line, human mean accuracy), the mean absolute error between predicted and measured accuracy (second column), the Pearson correlation between predicted and measured accuracy (third column) and the coefficient of determination with respect to the identity line (fourth column) are shown as a function of the decision-noise standard deviation *σ* (0, 0.25, 0.40–0.75 in steps of 0.05, and 1.0). Open circles mark the accuracy-matched *σ* of each condition used in the main analyses (0.60 for No CND and CND (LS+MS), 0.45 for CND (all fibers)). Axes are truncated for MAE and R^2^id at large *σ*, where accuracy falls far below the human range. For every read-out, the lowest MAE and the highest R^2^id reached by No CND or CND (LS+MS) at any noise level remained worse than those of CND of all fibers at its matched noise, and CND of all fibers gave the highest correlation with listeners at almost every noise level.

**Figure A8.**
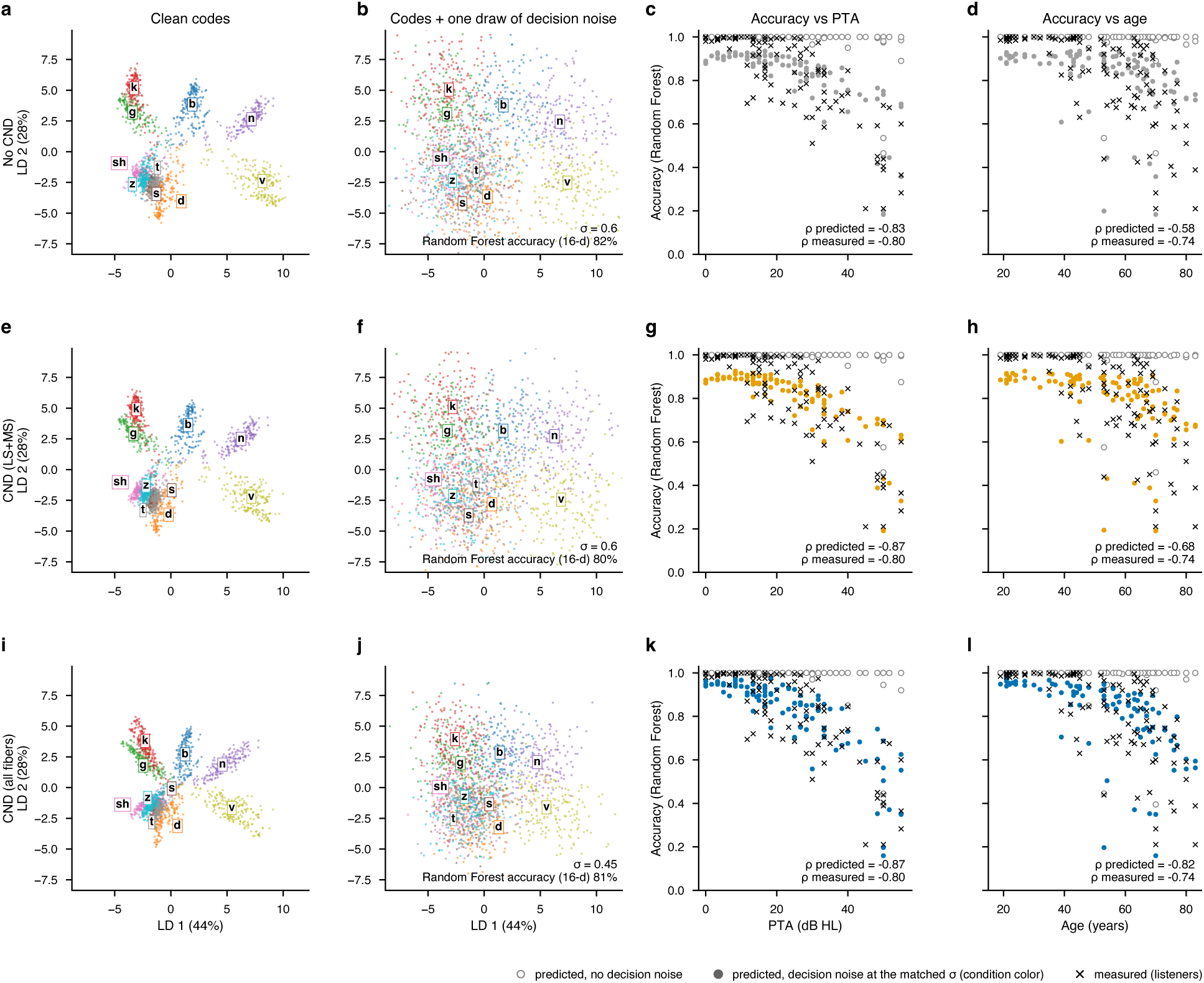
Decision noise lowers accuracy most for compressed codes; only with CND of all fibers do predictions follow listeners’ dependence on age and reach the ceiling of near-normal listeners. The learned codes with and without decision noise, and the resulting predictions against hearing loss and age, with each condition at its own accuracy-matched noise. Rows: audiometric loss only (No CND), with CND of low- and medium-spontaneous-rate fibers (CND (LS+MS)) and with CND of all fiber types (CND (all fibers)). Each condition is read out at its own accuracy-matched decision noise (*σ* = 0.60, 0.60 and 0.45), the level that brings its group-mean accuracy to the human mean and that is at or near the level giving its best fit to individual listeners (Fig. A7); each row therefore shows the most favorable setting of that condition. Fig. A9 shows the same analysis with a common noise level (*σ* = 0.45) for all conditions. (a, e, i) Clean 16-dimensional codes of all 94 listeners (20 tokens each) projected onto the two linear discriminants of Fig. 3, colored by consonant. (b, f, j) The same codes after adding one draw of the decision noise, projected with the same discriminants. The noise is added in all 16 dimensions, so this two-dimensional view, which shows only the two most discriminative directions, exaggerates the overlap between consonants; in the full code the random forest identifies 80–82% of tokens. (c, g, k) Accuracy of each listener against PTA: predicted by the random forest read-out without decision noise (open circles) and with decision noise (filled circles, condition color), and measured (crosses). (d, h, l) The same accuracies against age. Spearman *ρ* between PTA (or age) and predicted (with noise) or measured accuracy is given in each panel. Without noise, the read-out identifies nearly every token for every listener; the noise lowers accuracy most for listeners whose codes are compressed. All three conditions follow PTA about as closely as listeners do (*ρ* = 0.83 to 0.87 vs. 0.80). Without CND or with LS+MS CND, predictions depend less on age than measured accuracy does (*ρ* = 0.58 and 0.68 vs. 0.74), and listeners with near-normal hearing are predicted below their ceiling performance. With CND of all fibers, predictions match these listeners and follow age slightly more strongly than measured accuracy (*ρ* = 0.82). Codes in a, b, e, f, i and j are from the fold-1 network of Fig. 3; noise-free predictions in c, d, g, h, k and l are from networks trained without bottleneck noise.

**Figure A9.**
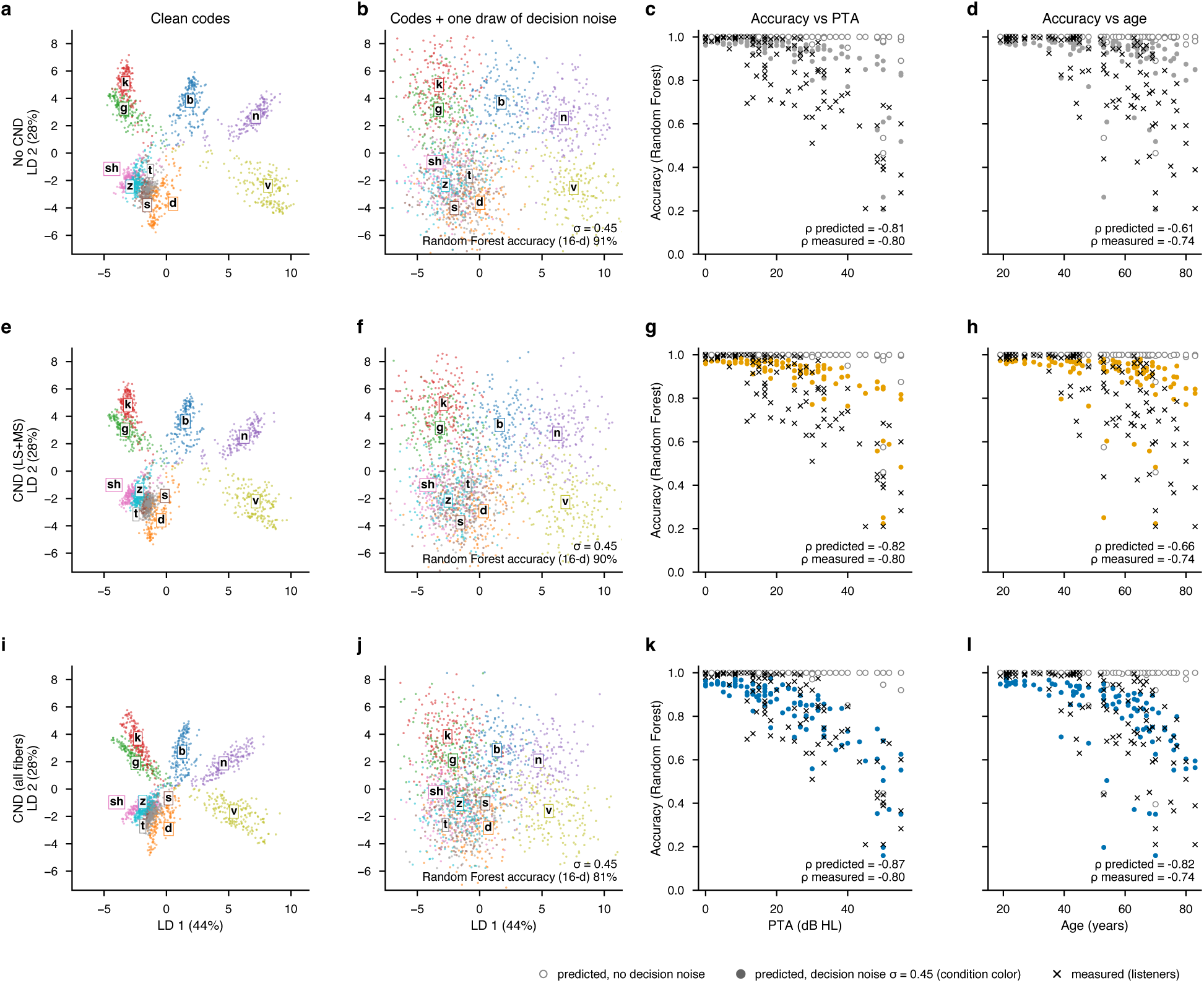
At the same decision noise, only CND of all fibers degrades the code enough to bring accuracy into the human range. Same as Fig. A8, with the same decision noise (*σ* = 0.45, the accuracy-matched level of CND of all fiber types) for all three conditions. Rows: audiometric loss only (No CND), with CND of low- and medium-spontaneous-rate fibers (CND (LS+MS)) and with CND of all fiber types (CND (all fibers)). (a, e, i) Clean 16-dimensional codes of all 94 listeners projected onto the two linear discriminants of Fig. 3, colored by consonant. (b, f, j) The same codes after adding one draw of decision noise with *σ* = 0.45, projected with the same discriminants; because the noise is added in all 16 dimensions, this view exaggerates the overlap between consonants. (c, g, k) Accuracy of each listener against PTA, predicted by the random forest read-out without decision noise (open circles) and with decision noise (filled circles, condition color), and measured (crosses). (d, h, l) The same accuracies against age. Spearman *ρ* between PTA (or age) and predicted (with noise) or measured accuracy is given in each panel. At this common noise level, the models without CND or with LS+MS CND overpredict accuracy (group means 91% and 90% vs. 82% for listeners), place most listeners near ceiling, and show a weaker dependence on age than measured accuracy (*ρ* = 0.61 and 0.66 vs. 0.74), whereas CND of all fibers matches the human accuracy range (81%). Thus, at the same decision noise, only degeneration across all fiber types degrades the simulated representation enough to reproduce listeners’ performance. Codes in a, b, e, f, i and j are from the fold-1 network of Fig. 3; noise-free predictions in c, d, g, h, k and l are from networks trained without bottleneck noise.

**Figure A10.**
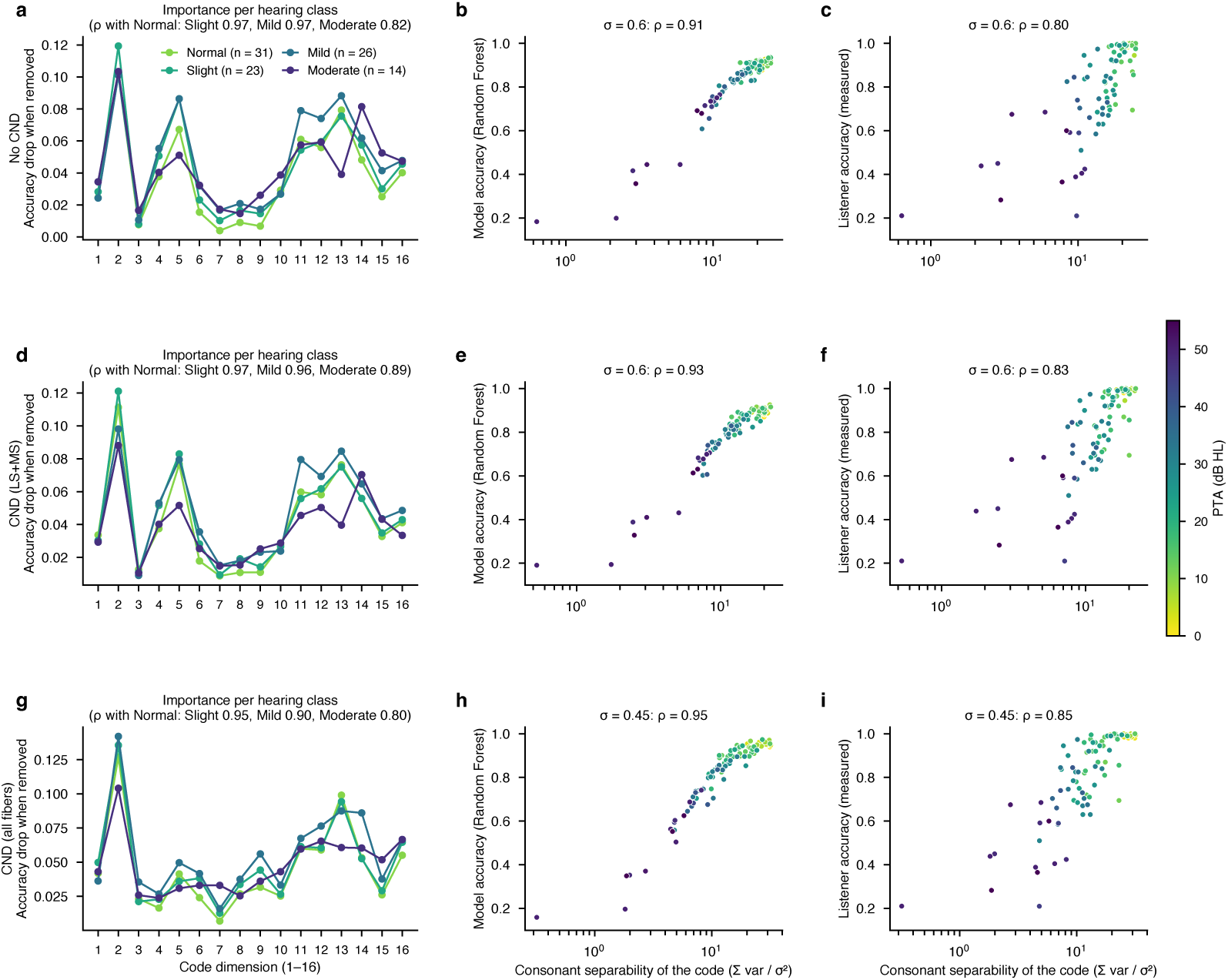
Hearing loss and CND reduce consonant separation along the same latent code dimensions in every hearing class, and this separation between phonemes tracks predicted and measured accuracy. Dimension importance and consonant separability of the learned code, computed for the held-out listeners of each cross-validation fold. Rows: audiometric loss only (No CND; a–c), with CND of low- and medium-spontaneous-rate fibers (CND (LS+MS); d–f) and with CND of all fiber types (CND (all fibers); g–i), each at its accuracy-matched decision noise (*σ* = 0.60, 0.60 and 0.45). (a, d, g) Which code dimensions the read-out relies on. In each fold, a random forest was trained on the codes of the training listeners (five noisy copies of each token) and applied to the codes of the held-out listeners with 20 draws of decision noise. Each of the 16 dimensions was then removed in turn by setting it to its mean over the training codes before the noise was added, and the importance of that dimension was the resulting drop in each listener’s accuracy. The same noise draws were used for the full and the reduced code, so the drop reflects only the removed dimension. Lines show the mean importance over the listeners of each hearing class (normal, slight, mild and moderate, as classified in the qVCV dataset), and titles give the Spearman rank correlation between the 16-value importance profile of normal-hearing listeners and that of each other class; a high correlation means that the same dimensions carry the consonant information for all classes. (b, e, h) How far apart each listener’s consonants lie in the code relative to the decision noise. For each listener, the noise-free codes of each consonant were averaged over its tokens, giving ten consonant means; in each dimension, the variance of these ten means was divided by *σ*^2^, and the result was summed over the 16 dimensions. This total separability is a signal-to-decision-noise ratio: the higher it is, the further apart the consonants lie compared with the spread added by the noise. It is computed from the simulated codes alone and is plotted against the model’s predicted accuracy (random forest, main read-out). (c, f, i) The same separability against the listener’s measured accuracy in the qVCV test. Points are listeners, colored by PTA; separability is shown on a logarithmic axis, and titles give the Spearman correlation across the 94 listeners. The same dimensions carry the consonant information in every condition and hearing class (rank correlations 0.80–0.97), so hearing loss and CND reduce separation along the dimensions that all listeners rely on rather than shifting information to other dimensions. Separability tracked predicted accuracy (Spearman *ρ* = 0.91, 0.93 and 0.95) and measured accuracy (*ρ* = 0.80, 0.83 and 0.85). The correlations are highest with CND of all fibers because age-scaled loss across all fiber types removes separation from the codes of older listeners in addition to the effect of their hearing loss: separability then varies more between listeners (coefficient of variation 0.55, vs. 0.36 and 0.38) and follows age as well as PTA (Spearman *ρ* with age = 0.88, vs. 0.67 and 0.77; with PTA 0.88 in all conditions), and so aligns more closely with listeners’ measured accuracy, which depends on both hearing loss and age.

## B Appendix tables

**Table A1.** Autoencoder architecture search. Each candidate architecture, differing in latent dimensionality and in the number and width of the convolutional encoder layers (the decoder mirrors the encoder), was trained exactly as the final model: a single autoencoder on the neurograms of all three conditions of the training listeners of each of the five cross-validation folds, with 15% of these listeners held out as an inner validation set for early stopping, and without noise in the latent code. All networks used 4 4 convolutions with stride 2, batch normalization and sigmoid activations, the Adam optimizer (learning rate 10*^−^*^3^, batch size 64) and early stopping (patience 12 epochs, at most 200 epochs). Entries are the latent dimensionality and means s.d. over the five folds of the mean squared reconstruction error on the inner-validation neurograms at the best epoch (Val MSE, 10*^−^*^4^; all three conditions pooled) and the reconstruction R^2^ of each condition, i.e. the proportion of the variance of the held-out listeners’ neurograms explained by their reconstructions. Reconstruction error fell steeply as the latent code grew from 2 to 16 dimensions and changed little among the four 16-dimensional architectures (a05–a08), which explained 92–96% of the held-out variance in every condition. The larger of these reduced the reconstruction error only slightly (by at most 0.7 10*^−^*^4^) at the cost of up to five times as many parameters, so we used the 16-dimensional architecture with the fewest parameters (a05, bold; 1.14 million). The layers of every candidate are listed in Table A3.

| **Arch.** | **Latent** | **Val MSE (×10<sup>−4</sup>)** | **R<sup>2</sup> No CND** | **R<sup>2</sup> CND (LS+MS)** | **R<sup>2</sup> CND (all fibers)** |
|---|---|---|---|---|---|
| a00 | 2 | 19.33 ± 0.74 | 0.587 ± 0.041 | 0.598 ± 0.042 | 0.718 ± 0.038 |
| a01 | 4 | 17.19 ± 0.70 | 0.639 ± 0.036 | 0.642 ± 0.038 | 0.753 ± 0.038 |
| a02 | 4 | 14.79 ± 3.41 | 0.692 ± 0.047 | 0.694 ± 0.048 | 0.785 ± 0.043 |
| a03 | 8 | 9.93 ± 1.85 | 0.788 ± 0.039 | 0.792 ± 0.039 | 0.861 ± 0.027 |
| a04 | 8 | 5.18 ± 0.70 | 0.895 ± 0.013 | 0.895 ± 0.014 | 0.931 ± 0.008 |
| **a05** | **16** | **3.47 ± 0.42** | **0.927 ± 0.012** | **0.925 ± 0.013** | **0.947 ± 0.011** |
| a06 | 16 | 3.18 ± 0.61 | 0.932 ± 0.018 | 0.931 ± 0.019 | 0.952 ± 0.013 |
| a07 | 16 | 3.06 ± 0.46 | 0.939 ± 0.003 | 0.937 ± 0.002 | 0.952 ± 0.004 |
| a08 | 16 | 2.81 ± 0.44 | 0.942 ± 0.008 | 0.940 ± 0.009 | 0.957 ± 0.009 |

**Table A2.**
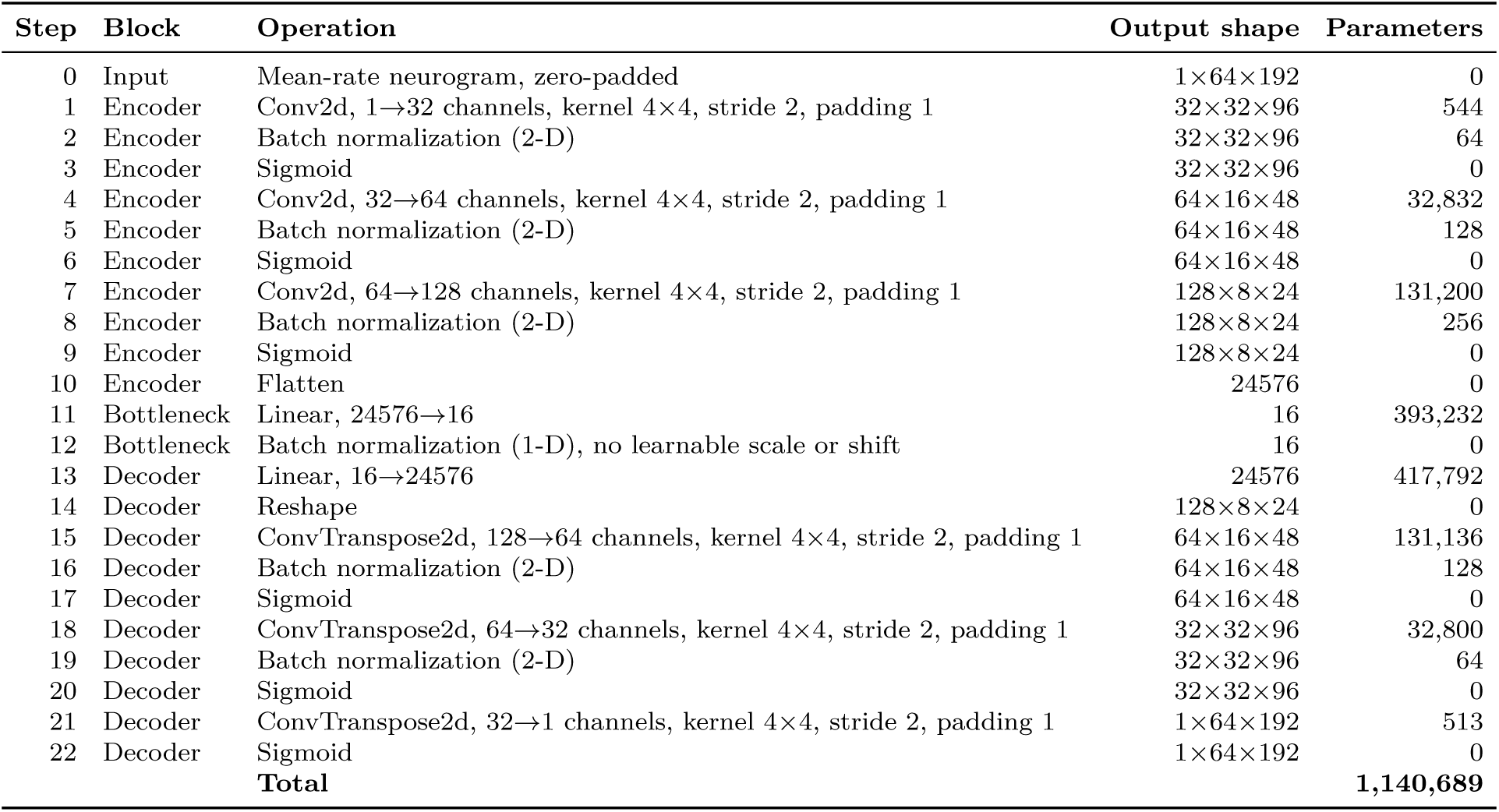
Layers of the selected autoencoder (a05). Every operation with its output shape (channels CF channels time bins) and number of trainable parameters, for an input neurogram zero-padded to 64 CF channels and 192 time bins (50 187 data region). The 16-dimensional latent code z (step 12) is batch-normalized without learnable scale or shift, so that each dimension has zero mean and unit variance; Gaussian noise is added to z before the decoder during training and before the classifiers during read-out. The loss is the mean squared error between input and reconstruction over the 50 × 187 data region.

**Table A3.**
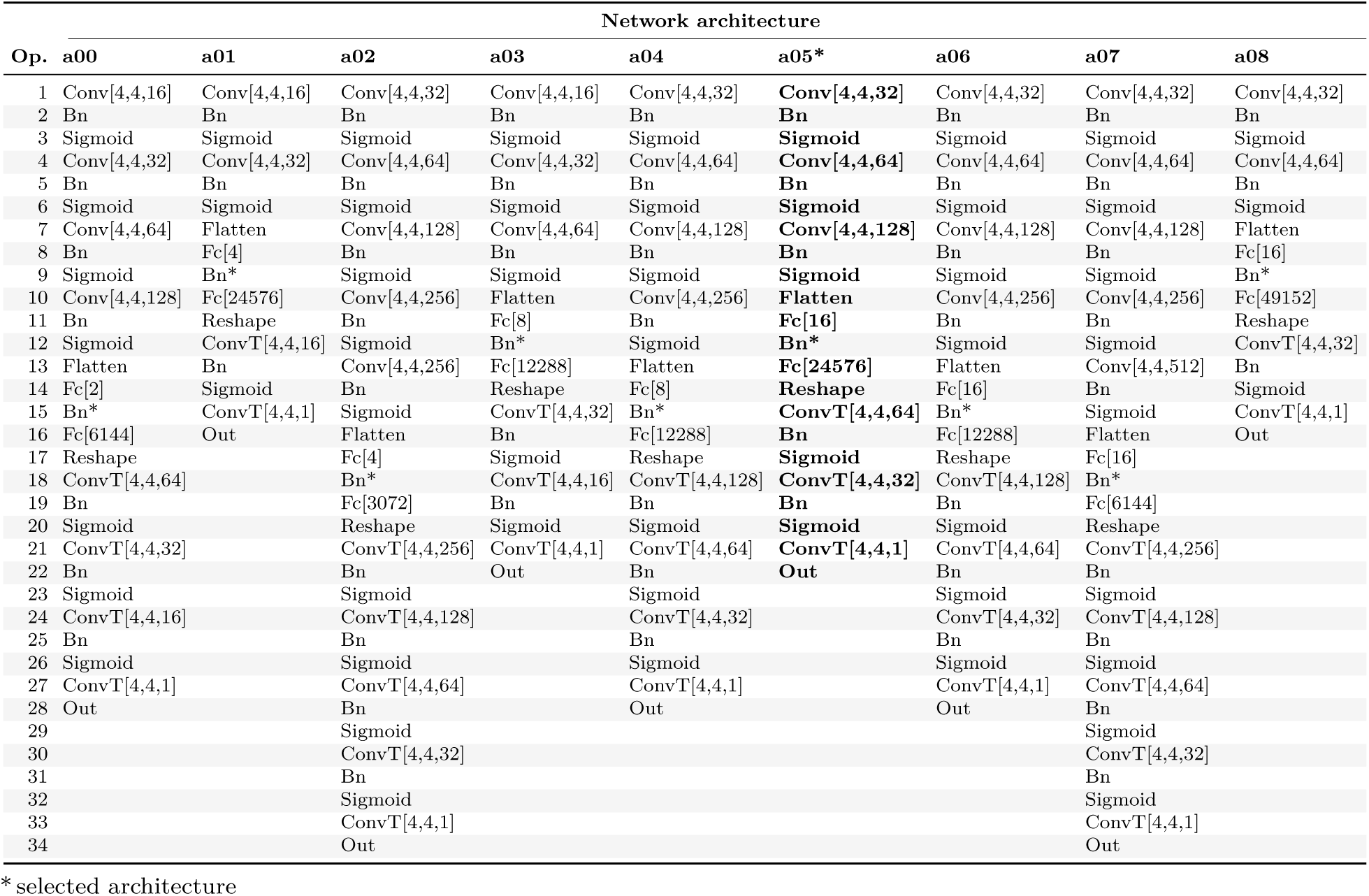
Operations of every candidate architecture of the search (Table A1), in order from input to reconstruction. All networks take a mean-rate neurogram zero-padded to 1 64 192 and are trained to reproduce it; the encoder halves the CF and time dimensions at each convolution, the bottleneck maps the result to the latent code (Fc[latent] followed by Bn*), and the decoder mirrors the encoder. a05 (bold) was used for all analyses. Layer keys are defined in the legend below.

|  | **Network architecture** |  |  |  |  |  |  |  |  |
|---|---|---|---|---|---|---|---|---|---|
| **Op.** | **a00** | **a01** | **a02** | **a03** | **a04** | **a05\*** | **a06** | **a07** | **a08** |
| 1 | Conv[4,4,16] | Conv[4,4,16] | Conv[4,4,32] | Conv[4,4,16] | Conv[4,4,32] | **Conv[4,4,32]** | Conv[4,4,32] | Conv[4,4,32] | Conv[4,4,32] |
| 2 | Bn | Bn | Bn | Bn | Bn | **Bn** | Bn | Bn | Bn |
| 3 | Sigmoid | Sigmoid | Sigmoid | Sigmoid | Sigmoid | **Sigmoid** | Sigmoid | Sigmoid | Sigmoid |
| 4 | Conv[4,4,32] | Conv[4,4,32] | Conv[4,4,64] | Conv[4,4,32] | Conv[4,4,64] | **Conv[4,4,64]** | Conv[4,4,64] | Conv[4,4,64] | Conv[4,4,64] |
| 5 | Bn | Bn | Bn | Bn | Bn | **Bn** | Bn | Bn | Bn |
| 6 | Sigmoid | Sigmoid | Sigmoid | Sigmoid | Sigmoid | **Sigmoid** | Sigmoid | Sigmoid | Sigmoid |
| 7 | Conv[4,4,64] | Flatten | Conv[4,4,128] | Conv[4,4,64] | Conv[4,4,128] | **Conv[4,4,128]** | Conv[4,4,128] | Conv[4,4,128] | Flatten |
| 8 | Bn | Fc[4] | Bn | Bn | Bn | **Bn** | Bn | Bn | Fc[16] |
| 9 | Sigmoid | Bn* | Sigmoid | Sigmoid | Sigmoid | **Sigmoid** | Sigmoid | Sigmoid | Bn* |
| 10 | Conv[4,4,128] | Fc[24576] | Conv[4,4,256] | Flatten | Conv[4,4,256] | **Flatten** | Conv[4,4,256] | Conv[4,4,256] | Fc[49152] |
| 11 | Bn | Reshape | Bn | Fc[8] | Bn | **Fc[16]** | Bn | Bn | Reshape |
| 12 | Sigmoid | ConvT[4,4,16] | Sigmoid | Bn* | Sigmoid | **Bn\*** | Sigmoid | Sigmoid | ConvT[4,4,32] |
| 13 | Flatten | Bn | Conv[4,4,256] | Fc[12288] | Flatten | **Fc[24576]** | Flatten | Conv[4,4,512] | Bn |
| 14 | Fc[2] | Sigmoid | Bn | Reshape | Fc[8] | **Reshape** | Fc[16] | Bn | Sigmoid |
| 15 | Bn* | ConvT[4,4,1] | Sigmoid | ConvT[4,4,32] | Bn* | **ConvT[4,4,64]** | Bn* | Sigmoid | ConvT[4,4,1] |
| 16 | Fc[6144] | Out | Flatten | Bn | Fc[12288] | **Bn** | Fc[12288] | Flatten | Out |
| 17 | Reshape |  | Fc[4] | Sigmoid | Reshape | **Sigmoid** | Reshape | Fc[16] |  |
| 18 | ConvT[4,4,64] |  | Bn* | ConvT[4,4,16] | ConvT[4,4,128] | **ConvT[4,4,32]** | ConvT[4,4,128] | Bn* |  |
| 19 | Bn |  | Fc[3072] | Bn | Bn | **Bn** | Bn | Fc[6144] |  |
| 20 | Sigmoid |  | Reshape | Sigmoid | Sigmoid | **Sigmoid** | Sigmoid | Reshape |  |
| 21 | ConvT[4,4,32] |  | ConvT[4,4,256] | ConvT[4,4,1] | ConvT[4,4,64] | **ConvT[4,4,1]** | ConvT[4,4,64] | ConvT[4,4,256] |  |
| 22 | Bn |  | Bn | Out | Bn | **Out** | Bn | Bn |  |
| 23 | Sigmoid |  | Sigmoid |  | Sigmoid |  | Sigmoid | Sigmoid |  |
| 24 | ConvT[4,4,16] |  | ConvT[4,4,128] |  | ConvT[4,4,32] |  | ConvT[4,4,32] | ConvT[4,4,128] |  |
| 25 | Bn |  | Bn |  | Bn |  | Bn | Bn |  |
| 26 | Sigmoid |  | Sigmoid |  | Sigmoid |  | Sigmoid | Sigmoid |  |
| 27 | ConvT[4,4,1] |  | ConvT[4,4,64] |  | ConvT[4,4,1] |  | ConvT[4,4,1] | ConvT[4,4,64] |  |
| 28 | Out |  | Bn |  | Out |  | Out | Bn |  |
| 29 |  |  | Sigmoid |  |  |  |  | Sigmoid |  |
| 30 |  |  | ConvT[4,4,32] |  |  |  |  | ConvT[4,4,32] |  |
| 31 |  |  | Bn |  |  |  |  | Bn |  |
| 32 |  |  | Sigmoid |  |  |  |  | Sigmoid |  |
| 33 |  |  | ConvT[4,4,1] |  |  |  |  | ConvT[4,4,1] |  |
| 34 |  |  | Out |  |  |  |  | Out |  |

<sup>\*</sup> selected architecture

### Layer legend

| **Key** | **Description** |
|---|---|
| Conv[X,Y,Z] | 2-D convolutional layer with kernel height X, kernel width Y and Z filters (stride 2, padding 1) |
| ConvT[X,Y,Z] | 2-D transposed convolutional layer with kernel height X, kernel width Y and Z filters (stride 2, padding 1) |
| Bn | Batch normalization layer |
| Bn* | Batch normalization without learnable scale or shift (latent code: zero mean and unit variance per dimension) |
| Sigmoid | Logistic sigmoid activation |
| Flatten | Reshape the feature maps into a vector |
| Fc[X] | Fully connected (linear) layer with X units; Fc[latent] defines the latent code |
| Reshape | Reshape the vector into feature maps (inverse of Flatten) |
| Out | Sigmoid output layer: reconstructed neurogram (1 × 64 × 192) |

**Table A4.**
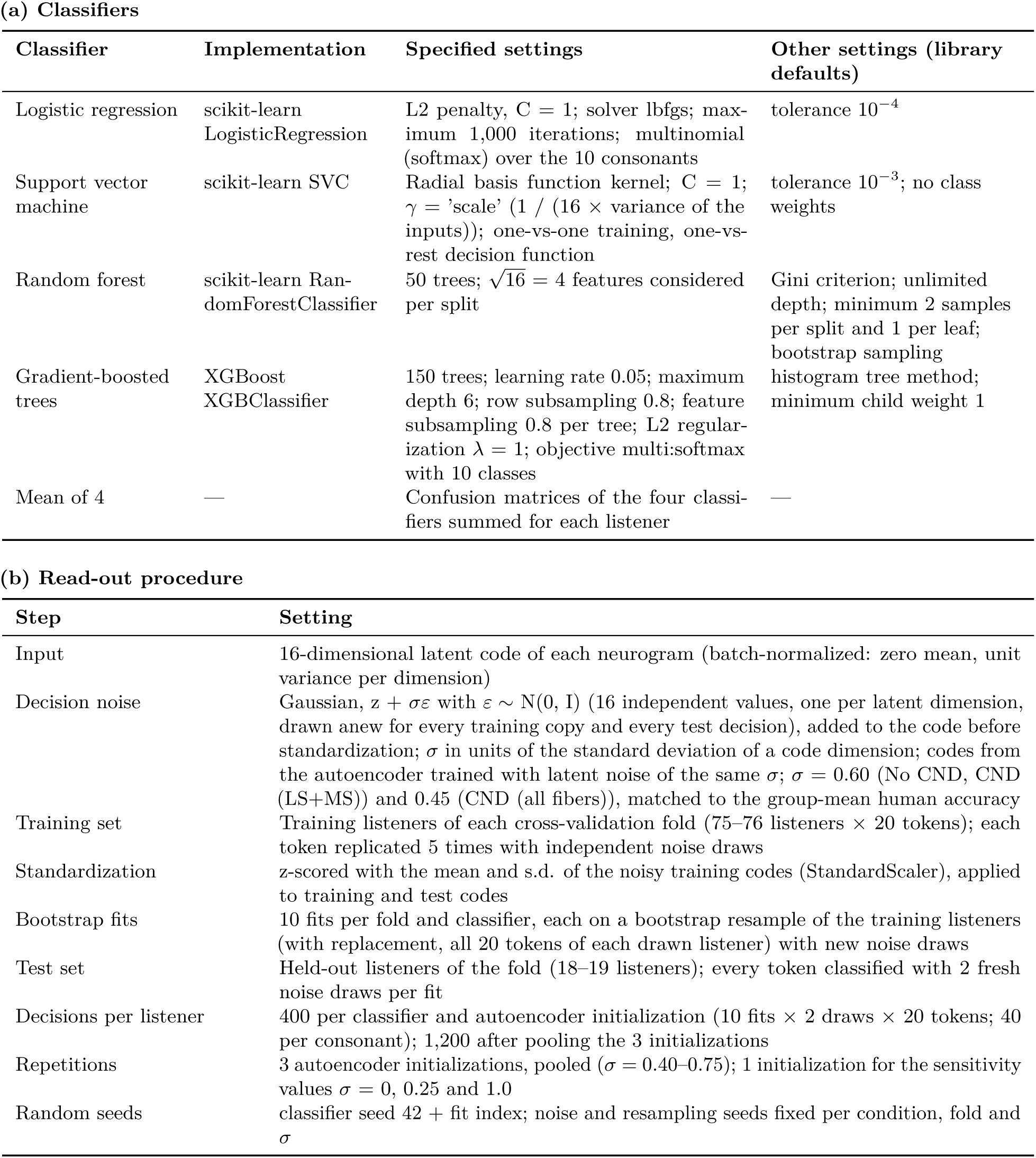
Downstream classifiers and read-out. (a) The four classifiers that identify the consonant from the 16-dimensional latent code, with the settings specified in the analysis code; all other settings were the library defaults (scikit-learn 1.7.2, XGBoost 3.2.0). Settings were fixed in advance and were not tuned. (b) The decision-noise read-out, identical for all classifiers and conditions.

| **(a) Classifiers** |  |  |  |
|---|---|---|---|
| **Classifier** | **Implementation** | **Specified settings** | **Other settings (library defaults)** |
| Logistic regression | scikit-learn LogisticRegression | L2 penalty, C = 1; solver lbfgs; max-imum 1,000 iterations; multinomial (softmax) over the 10 consonants | tolerance 10<sup>−4</sup> |
| Support vector machine | scikit-learn SVC | Radial basis function kernel; C = 1; γ = 'scale' (1 / (16 × variance of the inputs)); one-vs-one training, one-vs-rest decision function | tolerance 10<sup>−3</sup>; no class weights |
| Random forest | scikit-learn Ran-domForestClassifier | 50 trees; √16 = 4 features considered per split | Gini criterion; unlimited depth; minimum 2 samples per split and 1 per leaf; bootstrap sampling |
| Gradient-boosted trees | XGBoost XGBClassifier | 150 trees; learning rate 0.05; maximum depth 6; row subsampling 0.8; feature subsampling 0.8 per tree; L2 regular-ization λ = 1; objective multi:softmax with 10 classes | histogram tree method; minimum child weight 1 |
| Mean of 4 | — | Confusion matrices of the four classi-fiers summed for each listener | — |
| **(b) Read-out procedure** |  |  |  |
| **Step** | **Setting** |  |  |
| Input | 16-dimensional latent code of each neurogram (batch-normalized: zero mean, unit variance per dimension) |  |  |
| Decision noise | Gaussian, z + σε with ε ∼ N(0, I) (16 independent values, one per latent dimension, drawn anew for every training copy and every test decision), added to the code before standardization; σ in units of the standard deviation of a code dimension; codes from the autoencoder trained with latent noise of the same σ; σ = 0.60 (No CND, CND (LS+MS)) and 0.45 (CND (all fibers)), matched to the group-mean human accuracy |  |  |
| Training set | Training listeners of each cross-validation fold (75–76 listeners × 20 tokens); each token replicated 5 times with independent noise draws |  |  |
| Standardization | z-scored with the mean and s.d. of the noisy training codes (StandardScaler), applied to training and test codes |  |  |
| Bootstrap fits | 10 fits per fold and classifier, each on a bootstrap resample of the training listeners (with replacement, all 20 tokens of each drawn listener) with new noise draws |  |  |
| Test set | Held-out listeners of the fold (18–19 listeners); every token classified with 2 fresh noise draws per fit |  |  |
| Decisions per listener | 400 per classifier and autoencoder initialization (10 fits × 2 draws × 20 tokens; 40 per consonant); 1,200 after pooling the 3 initializations |  |  |
| Repetitions | 3 autoencoder initializations, pooled (σ = 0.40–0.75); 1 initialization for the sensitivity values σ = 0, 0.25 and 1.0 |  |  |
| Random seeds | classifier seed 42 + fit index; noise and resampling seeds fixed per condition, fold and σ |  |  |

## C Appendix algorithms

### Notation

Listeners *u* = 1,…, 94; tokens *t* = 1,…, 20 (10 consonants × 2 recordings); conditions *c*: No CND, CND (LS+MS) and CND (all); cross-validation folds *k* = 1,…, 5 (PTA-stratified, 18–19 held-out listeners each); ***Y****_k_* and *V_k_* are the training and held-out listeners of fold *k*; *X_u,t,c_ ∈* ℝ^50^*^×^*^187^ is the mean-rate neurogram of listener *u*, token *t* and condition *c*; *N*(0*, I*) is the standard normal distribution in the 16-dimensional latent space.

#### Algorithm 1

Training of the shared autoencoder for fold *k*, initialization seed *r* and noise level *σ*

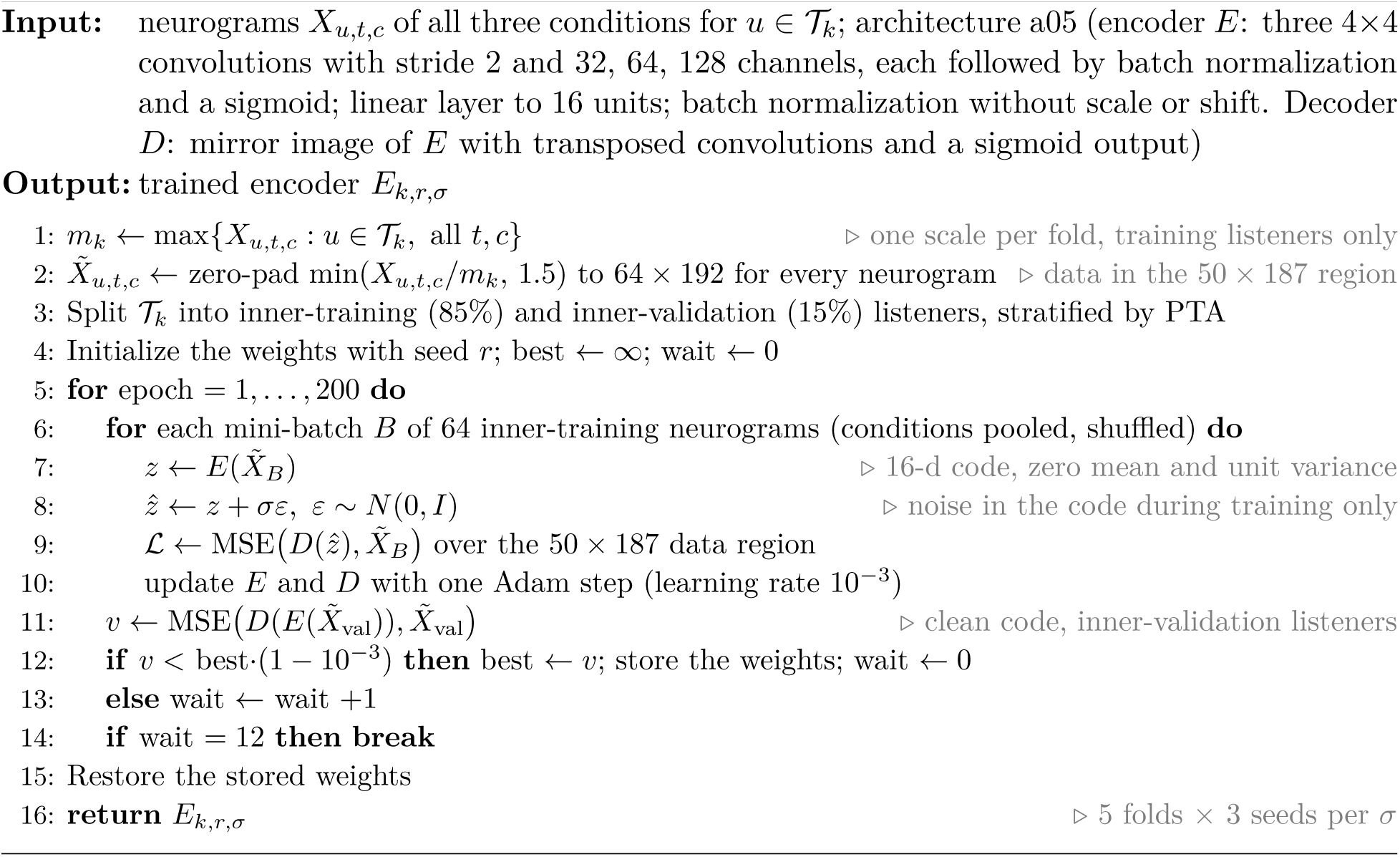

#### Algorithm 2

Training of the downstream classifiers and decision-noise read-out for fold *k*

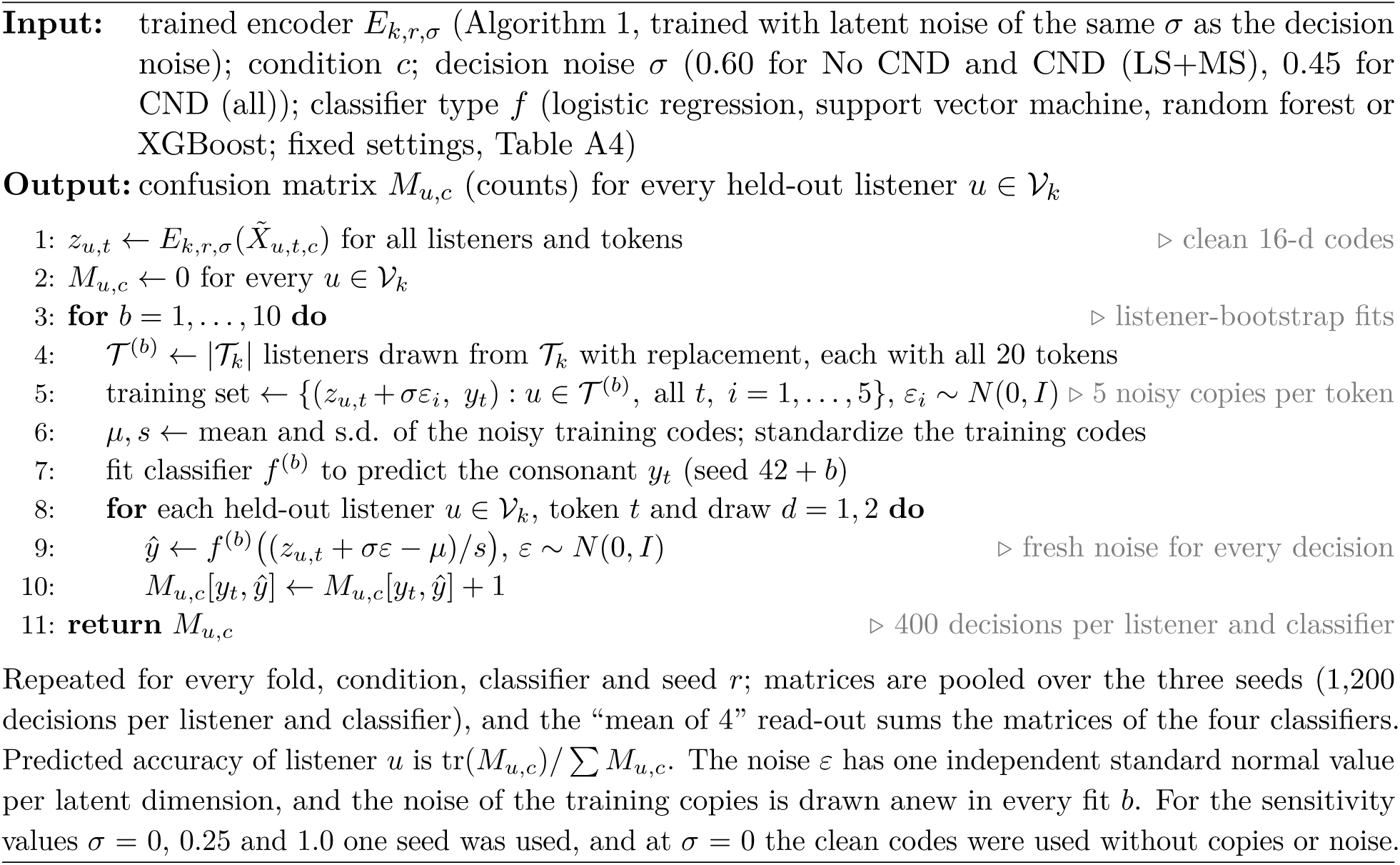

## Notes

### Competing Interest Statement

The authors have declared no competing interest.

## References

[1] Kelsie J. Grant, Aravindakshan Parthasarathy, Viacheslav Vasilkov, Benjamin Caswell-Midwinter, Maria E. Freitas, Victor de Gruttola, Daniel B. Polley, M. Charles Liberman, and Stéphane F. Maison. Predicting neural deficits in sensorineural hearing loss from word recognition scores. Scientific Reports, 12(1):8929, 2022. doi: 10.1038/s41598-022-13023-5.

[2] Martin Pienkowski. On the etiology of listening difficulties in noise despite clinically normal audiograms. Ear & Hearing, 38(2):135–148, 2017. doi: 10.1097/aud.0000000000000388.

[3] A. C. Carl and J. Cornejo. Audiology pure tone evaluation. In StatPearls. StatPearls Publishing, Treasure Island (FL), 2022. URL https://www.ncbi.nlm.nih.gov/books/NBK580531/. NCBI Bookshelf NBK580531.

[4] Seung Yeol Lee, Hee Won Seo, Seon Min Jung, Seung Hwan Lee, and Jae Ho Chung. Assessing the accuracy and reliability of application-based audiometry for hearing evaluation. Scientific Reports, 14 (1):7359, 2024. doi: 10.1038/s41598-024-57944-9.

[5] Curtis J. Billings, Tessa M. Olsen, Lauren Charney, Brandon M. Madsen, and Corrie E. Holmes. Speech-in-noise testing: An introduction for audiologists. Seminars in Hearing, 45(01):055–082, 2023. doi: 10.1055/s-0043-1770155.

[6] Lidia W. Lee and Larry E. Humes. Evaluating a speech-reception threshold model for hearing-impaired listeners. The Journal of the Acoustical Society of America, 93(5):2879–2885, 1993. doi: 10.1121/1.405807.

[7] Judy R. Dubno, Fu-Shing Lee, Arnold J. Klein, Lois J. Matthews, and Chin Fai Lam. Confidence limits for maximum word-recognition scores. Journal of Speech and Hearing Research, 38(2):490–502, 1995. doi: 10.1044/jshr.3802.490.

[8] Chris Halpin and Steven D. Rauch. Clinical implications of a damaged cochlea: Pure tone thresholds vs information-carrying capacity. Otolaryngology–Head and Neck Surgery, 140(4):473–476, 2009. doi: 10.1016/j.otohns.2008.12.021.

[9] Anita M. Mepani, Sarah Verhulst, Kenneth E. Hancock, Markus Garrett, Viacheslav Vasilkov, Kara Bennett, Victor de Gruttola, M. Charles Liberman, and Stéphane F. Maison. Envelope following responses predict speech-in-noise performance in normal-hearing listeners. Journal of Neurophysiology, 125(4):1213–1222, 2021. doi: 10.1152/jn.00620.2020.

[10] Jiayue Liu, Joshua Stohl, and Tobias Overath. Hidden hearing loss: Fifteen years at a glance. Hearing Research, 443:108967, 2024. doi: 10.1016/j.heares.2024.108967.

[11] Richard H. Wilson. Clinical experience with the words-in-noise test on 3430 veterans: Comparisons with pure-tone thresholds and word recognition in quiet. Journal of the American Academy of Audiology, 22 (7):405–423, 2011. doi: 10.3766/jaaa.22.7.3.

[12] Luis Robles and Mario A. Ruggero. Mechanics of the mammalian cochlea. Physiological Reviews, 81(3): 1305–1352, 2001. doi: 10.1152/physrev.2001.81.3.1305.

[13] P Dallos. The active cochlea. The Journal of Neuroscience, 12(12):4575–4585, 1992. doi: 10.1523/jneurosci.12-12-04575.1992.

[14] Brian C.J. Moore. Cochlear Hearing Loss: Physiological, Psychological and Technical Issues. Wiley, 2nd edition, 2007. doi: 10.1002/9780470987889.

[15] Larry E. Humes, Judy R. Dubno, Sandra Gordon-Salant, Jennifer J. Lister, Anthony T. Cacace, Karen J. Cruickshanks, George A. Gates, Richard H. Wilson, and Arthur Wingfield. Central presbycusis: A review and evaluation of the evidence. Journal of the American Academy of Audiology, 23(8):635–666, 2012. doi: 10.3766/jaaa.23.8.5.

[16] Brian C. J. Moore. Effects of age and hearing loss on the processing of auditory temporal fine structure. In Physiology, Psychoacoustics and Cognition in Normal and Impaired Hearing, pages 1–8. Springer International Publishing, 2016. doi: 10.1007/978-3-319-25474-6_1.

[17] Samira Anderson. Hearing and aging effects on speech understanding: Challenges and solutions. Acoustics Today, 14(4):10, 2018. doi 10.1121/at.2018.14.4.12.

[18] Larry E. Humes and Judy R. Dubno. Factors affecting speech understanding in older adults. In The Aging Auditory System, pages 211–257. Springer New York, 2009. doi: 10.1007/978-1-4419-0993-0_8.

[19] M. Charles Liberman and Sharon G. Kujawa. Cochlear synaptopathy in acquired sensorineural hearing loss: Manifestations and mechanisms. Hearing Research, 349:138–147, 2017. doi: 10.1016/j.heares.2017.01.003.

[20] Sharon G. Kujawa and M. Charles Liberman. Adding insult to injury: Cochlear nerve degeneration after “temporary” noise-induced hearing loss. The Journal of Neuroscience, 29(45):14077–14085, 2009. doi: 10.1523/jneurosci.2845-09.2009.

[21] Sharon G. Kujawa and M. Charles Liberman. Synaptopathy in the noise-exposed and aging cochlea: Primary neural degeneration in acquired sensorineural hearing loss. Hearing Research, 330:191–199, 2015. doi: 10.1016/j.heares.2015.02.009.

[22] Edward Lobarinas, Richard Salvi, and Dalian Ding. Insensitivity of the audiogram to carboplatin induced inner hair cell loss in chinchillas. Hearing Research, 302:113–120, 2013. doi: 10.1016/j.heares.2013.03.012.

[23] Lucas M. Viana, Jennifer T. O’Malley, Barbara J. Burgess, Dianne D. Jones, Carlos A.C.P. Oliveira, Felipe Santos, Saumil N. Merchant, Leslie D. Liberman, and M. Charles Liberman. Cochlear neuropathy in human presbycusis: Confocal analysis of hidden hearing loss in post-mortem tissue. Hearing Research, 327:78–88, 2015. doi: 10.1016/j.heares.2015.04.014.

[24] P.Z. Wu, L.D. Liberman, K. Bennett, V. de Gruttola, J.T. O’Malley, and M.C. Liberman. Primary neural degeneration in the human cochlea: Evidence for hidden hearing loss in the aging ear. Neuroscience, 407:8–20, 2019. doi: 10.1016/j.neuroscience.2018.07.053.

[25] Pei-Zhe Wu, Jennifer T. O’Malley, Victor de Gruttola, and M. Charles Liberman. Primary neural degeneration in noise-exposed human cochleas: Correlations with outer hair cell loss and word-discrimination scores. The Journal of Neuroscience, 41(20):4439–4447, 2021. doi: 10.1523/jneurosci.3238-20.2021.

[26] Hari M. Bharadwaj, Sarah Verhulst, Luke Shaheen, M. Charles Liberman, and Barbara G. Shinn-Cunningham. Cochlear neuropathy and the coding of supra-threshold sound. Frontiers in Systems Neuroscience, 8, 2014. doi: 10.3389/fnsys.2014.00026.

[27] Adam C. Furman, Sharon G. Kujawa, and M. Charles Liberman. Noise-induced cochlear neuropathy is selective for fibers with low spontaneous rates. Journal of Neurophysiology, 110(3):577–586, 2013. doi: 10.1152/jn.00164.2013.

[28] Christopher J. Plack, Daphne Barker, and Garreth Prendergast. Perceptual consequences of “hidden” hearing loss. Trends in Hearing, 18:2331216514550621, 2014. doi: 10.1177/2331216514550621.

[29] Naomi F. Bramhall, Dawn Konrad-Martin, Garnett P. McMillan, and Susan E. Griest. Auditory brainstem response altered in humans with noise exposure despite normal outer hair cell function. Ear & Hearing, 38(1):e1–e12, 2017. doi: 10.1097/aud.0000000000000370.

[30] Roland Schaette and David McAlpine. Tinnitus with a normal audiogram: Physiological evidence for hidden hearing loss and computational model. The Journal of Neuroscience, 31(38):13452–13457, 2011. doi: 10.1523/jneurosci.2156-11.2011.

[31] Hari M. Bharadwaj, Salwa Masud, Golbarg Mehraei, Sarah Verhulst, and Barbara G. Shinn-Cunningham. Individual differences reveal correlates of hidden hearing deficits. The Journal of Neuroscience, 35(5): 2161–2172, 2015. doi: 10.1523/jneurosci.3915-14.2015.

[32] Luke A. Shaheen, Michelle D. Valero, and M. Charles Liberman. Towards a diagnosis of cochlear neuropathy with envelope following responses. Journal of the Association for Research in Otolaryngology, 16(6):727–745, 2015. doi: 10.1007/s10162-015-0539-3.

[33] Viacheslav Vasilkov, Markus Garrett, Manfred Mauermann, and Sarah Verhulst. Enhancing the sensitivity of the envelope-following response for cochlear synaptopathy screening in humans: The role of stimulus envelope. Hearing Research, 400:108132, 2021. doi: 10.1016/j.heares.2020.108132.

[34] Anita M. Mepani, Sarah A. Kirk, Kenneth E. Hancock, Kara Bennett, Victor de Gruttola, M. Charles Liberman, and Stéphane F. Maison. Middle ear muscle reflex and word recognition in “normal-hearing” adults: Evidence for cochlear synaptopathy? Ear & Hearing, 41(1):25–38, 2019. doi: 10.1097/aud.0000000000000804.

[35] Michelle D. Valero, Kenneth E. Hancock, and M. Charles Liberman. The middle ear muscle reflex in the diagnosis of cochlear neuropathy. Hearing Research, 332:29–38, 2016. doi: 10.1016/j.heares.2015.11.005.

[36] Michelle D. Valero, Kenneth E. Hancock, Stéphane F. Maison, and M. Charles Liberman. Effects of cochlear synaptopathy on middle-ear muscle reflexes in unanesthetized mice. Hearing Research, 363: 109–118, 2018. doi: 10.1016/j.heares.2018.03.012.

[37] Kai Burian, Ahsan J. Cheema, and Sunil Puria. Modeling loudness contours to determine human auditory nerve fiber distribution and cochlear nerve degeneration. Proceedings of Meetings on Acoustics, 60(1):050006, 2025. doi 10.1121/2.0002265. 189th Meeting of the Acoustical Society of America and the Acoustical Society of Japan.

[38] Ahsan Jamal Cheema and Sunil Puria. Utilizing information theoretic approach to study cochlear neural degeneration. In ICASSP 2026 - 2026 IEEE International Conference on Acoustics, Speech and Signal Processing (ICASSP), pages 15562–15566. IEEE, 2026. doi: 10.1109/icassp55912.2026.11460958.

[39] Ahsan J. Cheema, David Meng, Jorge Mejia, Sanna Hou, and Sunil Puria. Model-guided design of low-context speech probes for cochlear synaptopathy. arXiv preprint arXiv:2609.36272, 2026.

[40] Naomi Bramhall, Elizabeth Francis Beach, Bastian Epp, Colleen G. Le Prell, Enrique A. Lopez-Poveda, Christopher J. Plack, Roland Schaette, Sarah Verhulst, and Barbara Canlon. The search for noise-induced cochlear synaptopathy in humans: Mission impossible? Hearing Research, 377:88–103, 2019. doi: 10.1016/j.heares.2019.02.016.

[41] Hannah Guest, Kevin J. Munro, Garreth Prendergast, Rebecca E. Millman, and Christopher J. Plack. Impaired speech perception in noise with a normal audiogram: No evidence for cochlear synaptopathy and no relation to lifetime noise exposure. Hearing Research, 364:142–151, 2018. doi: 10.1016/j.heares.2018.03.008.

[42] Garreth Prendergast, Hannah Guest, Kevin J. Munro, Karolina Kluk, Agnès Léger, Deborah A. Hall, Michael G. Heinz, and Christopher J. Plack. Effects of noise exposure on young adults with normal audiograms i: Electrophysiology. Hearing Research, 344:68–81, 2017. doi: 10.1016/j.heares.2016.10.028.

[43] Peter T. Johannesen and Enrique A. Lopez-Poveda. Age-related central gain compensation for reduced auditory nerve output for people with normal audiograms, with and without tinnitus. iScience, 24(6): 102658, 2021. doi 10.1016/j.isci.2021.102658.

[44] Feifan Chen, Fei Zhao, Nadeem Mahafza, and Wei Lu. Detecting noise-induced cochlear synaptopathy by auditory brainstem response in tinnitus patients with normal hearing thresholds: A meta-analysis. Frontiers in Neuroscience, 15:778197, 2021. doi: 10.3389/fnins.2021.778197.

[45] Kelsie J. Grant, Anita M. Mepani, Peizhe Wu, Kenneth E. Hancock, Victor de Gruttola, M. Charles Liberman, and Stéphane F. Maison. Electrophysiological markers of cochlear function correlate with hearing-in-noise performance among audiometrically normal subjects. Journal of Neurophysiology, 124 (2):418–431, 2020. doi: 10.1152/jn.00016.2020.

[46] Pei-zhe Wu, Jennifer T. O’Malley, and M. Charles Liberman. Neural degeneration in normal-aging human cochleas: Machine-learning counts and 3D mapping in archival sections. Journal of the Association for Research in Otolaryngology, 24(5):499–511, 2023. doi: 10.1007/s10162-023-00909-y.

[47] Aryn M. Kamerer, Sara E. Harris, Chris S. Wichman, Daniel M. Rasetshwane, and Stephen T. Neely. The relationship and interdependence of auditory thresholds, proposed behavioural measures of hidden hearing loss, and physiological measures of auditory function. International Journal of Audiology, 64(1): 11–24, 2024. doi: 10.1080/14992027.2024.2391986.

[48] Mishaela DiNino, Lori L. Holt, and Barbara G. Shinn-Cunningham. Cutting through the noise: Noise-induced cochlear synaptopathy and individual differences in speech understanding among listeners with normal audiograms. Ear & Hearing, 43(1):9–22, 2021. doi: 10.1097/aud.0000000000001147.

[49] Kanae Nishi, Andrea C. Trevino, Lydia Rosado Rogers, Paula García, and Stephen T. Neely. Effects of simulated hearing loss on bilingual children’s consonant recognition in noise. Ear & Hearing, 38(5): e292–e304, 2017. doi: 10.1097/aud.0000000000000428.

[50] Joshua J. Hajicek, Sara E. Harris, and Stephen Neely. Using consonant confusion to predict unexplained hearing loss. The Journal of the Acoustical Society of America, 154(4_supplement):A34–A34, 2023. doi: 10.1121/10.0022707.

[51] Joshua J. Hajicek, Sara E. Harris, and Stephen T. Neely. Development and clinical application of a consonant confusion task to evaluate hearing aid benefit. medRxiv, 2026. doi: 10.64898/2026.04.23.26351598. URL https://www.medrxiv.org/content/early/2026/04/24/2026.04.23.26351598. preprint 2026.04.23.26351598.

[52] Ahsan Jamal Cheema and Sunil Puria. Using neurogram similarity index measure (NSIM) to model hearing loss and cochlear neural degeneration. In Proc. Interspeech 2025, pages 823–827, 2025. doi: 10.21437/Interspeech.2025-2458.

[53] Sarah Verhulst, Alessandro Altoè, and Viacheslav Vasilkov. Computational modeling of the human auditory periphery: Auditory-nerve responses, evoked potentials and hearing loss. Hearing Research, 360:55–75, 2018. doi: 10.1016/j.heares.2017.12.018.

[54] Sarineh Keshishzadeh, Markus Garrett, and Sarah Verhulst. Towards personalized auditory models: Predicting individual sensorineural hearing-loss profiles from recorded human auditory physiology. Trends in Hearing, 25:2331216520988406, 2021. doi: 10.1177/2331216520988406.

[55] Markus Garrett, Viacheslav Vasilkov, Manfred Mauermann, Pauline Devolder, John L. Wilson, Leslie Gonzales, Kenneth S. Henry, and Sarah Verhulst. Deciphering compromised speech-in-noise intelligibility in older listeners: The role of cochlear synaptopathy. eneuro, 12(2):ENEURO.0182–24.2024, 2025. doi: 10.1523/eneuro.0182-24.2024.

[56] Enrique A. Lopez-Poveda and Pablo Barrios. Perception of stochastically undersampled sound waveforms: a model of auditory deafferentation. Frontiers in Neuroscience, 7, 2013. doi: 10.3389/fnins.2013.00124.

[57] Jiayue Liu, Joshua Stohl, Enrique A. Lopez-Poveda, and Tobias Overath. Quantifying the impact of auditory deafferentation on speech perception. Trends in Hearing, 28:23312165241227818, 2024. doi: 10.1177/23312165241227818.

[58] Johannes Zaar and Laurel H. Carney. Predicting speech intelligibility in hearing-impaired listeners using a physiologically inspired auditory model. Hearing Research, 426:108553, 2022. doi: 10.1016/j.heares.2022.108553.

[59] Fotios Drakopoulos, Viacheslav Vasilkov, Alejandro Osses Vecchi, Tijmen Wartenberg, and Sarah Verhulst. Model-based hearing-enhancement strategies for cochlear synaptopathy pathologies. Hearing Research, 424:108569, 2022. doi: 10.1016/j.heares.2022.108569.

[60] Fotios Drakopoulos and Sarah Verhulst. A neural-network framework for the design of individualised hearing-loss compensation. IEEE/ACM Transactions on Audio, Speech, and Language Processing, 31: 2395–2409, 2023. doi: 10.1109/taslp.2023.3282093.

[61] Mark R. Saddler and Josh H. McDermott. Models optimized for real-world tasks reveal the task-dependent necessity of precise temporal coding in hearing. Nature Communications, 15(1):10590, 2024. doi 10.1038/s41467-024-54700-5.

[62] Mark R. Saddler, Torsten Dau, and Josh H. McDermott. Towards individualized models of hearing-impaired speech perception. In The 6th Clarity Workshop on Improving Speech-in-Noise for Hearing Devices (Clarity-2025), pages 7–11. ISCA, 2025. doi: 10.21437/clarity.2025-3.

[63] Viacheslav Vasilkov, M. Charles Liberman, and Stéphane F. Maison. Modeling the impact of cochlear nerve degeneration on speech recognition performance. PLOS One, 20(11):e0336299, 2025. doi 10.1371/journal.pone.0336299.

[64] Annesya Banerjee, Mark R. Saddler, Julie G. Arenberg, and Josh H. McDermott. A deep learning framework for understanding cochlear implants. bioRxiv, 2025. doi: 10.1101/2025.07.16.665227. Preprint.

[65] Ian C. Bruce, Abigail Buller, and Muhammad S. Zilany. Modeling of auditory nerve fiber input/output functions near threshold. The Journal of the Acoustical Society of America, 154(4_supplement):A240–A240, 2023. doi: 10.1121/10.0023407.

[66] Ian C. Bruce, Yousof Erfani, and Muhammad S.A. Zilany. A phenomenological model of the synapse between the inner hair cell and auditory nerve: Implications of limited neurotransmitter release sites. Hearing Research, 360:40–54, 2018. doi: 10.1016/j.heares.2017.12.016.

[67] Jahangir Moini, Anthony LoGalbo, and Raheleh Ahangari. The auditory system. In Foundations of the Mind, Brain, and Behavioral Relationships, chapter 9, pages 143–160. Academic Press, 2024. ISBN 9780323959759. doi: 10.1016/B978-0-323-95975-9.00004-4.

[68] Brian C.J. Moore and Brian R. Glasberg. A revised model of loudness perception applied to cochlear hearing loss. Hearing Research, 188(1-2):70–88, 2004. doi: 10.1016/s0378-5955(03)00347-2.

[69] Donald D. Greenwood. A cochlear frequency-position function for several species—29 years later. The Journal of the Acoustical Society of America, 87(6):2592–2605, 1990. doi: 10.1121/1.399052.

[70] R. A. Schmiedt, J. H. Mills, and F. A. Boettcher. Age-related loss of activity of auditory-nerve fibers. Journal of Neurophysiology, 76(4):2799–2803, 1996. doi: 10.1152/jn.1996.76.4.2799.

